# High-Throughput De Novo Protein Design Yields Novel Immunomodulatory Agonists

**DOI:** 10.1101/2025.10.12.681920

**Authors:** Mohamad Abedi, Marc Expòsit, Brian Coventry, Divij Mathew, Shruti Jain, Aditya Krishnakumar, Inna Goreshnik, Sophie L. Gray-Gaillard, Margaret Lunn-Halbert, Ta-Yi Yu, Matthias Glögl, Uma Mitchell, Riya Keshri, Jung Ho Chun, Hannele Ruohola-Baker, E. John Wherry, David Baker

## Abstract

Cytokines regulate cell behavior by bringing together specific receptor subunits to trigger downstream signaling. Designed molecules that bring together non-natural receptor pairs could have novel signaling responses and cell specificities. We present a high-throughput de-novo design approach to create novel cytokines by generating and fusing pairs of computationally designed binders. By combining 33 designed receptor-binding domains, we generated over a thousand potential de novo designed “Novokines”, of which 75 activated pSTAT signaling in peripheral blood mononuclear cells. We characterized 18 of these, including new pairings of established common receptors, cross-family pairings such as TrkA-γcommon, and a series of pairings with interferon receptor-1 (IFNAR1), revealing that IFNAR1 can function as a versatile common receptor similar to γcommon or βcommon. We identify novokines that drive monocyte proliferation, T cell survival and CD4^+^ T cell-specific proliferation. Our framework provides a blueprint for expanding the understanding of cytokine signaling and generating novel therapeutic proteins.

## Introduction

Cytokines and growth factors exert their effects by inducing homo- or heterodimerization of receptor subunits, which initiates transphosphorylation of intracellular kinase domains and activates downstream signaling cascades. These receptor chains are organized into “common” subunits, which are shared across multiple cytokines to integrate and coordinate signaling (e.g., γcommon or βcommon), and “private” subunits, which are unique to individual ligands and confer specificity of recognition and response. To date, nearly all studies of these signaling pathways have focused on responses to naturally occurring ligands^1–3^, limiting our understanding of whether signaling can be triggered by receptor pairings that do not occur in nature. Receptor pairs have been dimerized using synthetic protein dimerizers in cell lines, but such systems are less reflective of natural cellular responses^4^. Synthetic ligands (synthekines) generated by linking nanobodies which bring together non-natural receptor pairs have demonstrated the potential to induce cellular responses in primary cells^5–7^, but only a few synthekines have been reported. Recent work expanded this potential by engineering primary T cells with novel receptor pairs responsive to orthogonal ligands, yielding signaling outcomes unattainable with natural cytokines and even redirecting T cells toward monocyte-like fates. While suitable for adoptive cell therapy applications, the requirement for engineered cells limits applications of this strategy for protein therapeutics based approaches^8^. Large-scale testing and characterization of agonists that can act on cells in their native context could greatly expand our understanding of cell signaling and lead to new therapeutic strategies. Computational protein design has been used to generate receptor agonists, but efforts have so far focused on a small number of natural receptor pairs (e.g., Erythropoietin receptor, IL-2, IL-4, IL-21)^9–12^.

We reasoned that recent advances in protein design, gene synthesis, and massively parallel protein expression could enable the systematic design and characterization of thousands of de novo designed signaling molecules (we refer to these as Novokines throughout the text) that bring receptor subunits into close proximity to probe their signaling effects. To do this, we set out to design binders targeting individual receptor subunits, fuse them using flexible or rigid linkers, either to one another or to previously developed binding domains, and assess their signaling activity in cells.

## Results

### De novo design of receptor binding proteins

As a first step towards generating a wide range of potential signaling molecules, we designed binders targeting a diverse set of cytokine and growth factor receptor subunits. Our selection was guided by two main objectives: modulating immune cell signaling, particularly in T cells, and enabling cross-family pairing of receptor subunits. We included cytokine receptors essential for immune function, such as IL4Rα, IL2Rβ, IL12Rβ1, IL10Rβ, IFNAR1, IFNAR2, and TSLPR, along with shared signaling subunits like γcommon (CD132) and βcommon (CD131).

To generate these binders, we employed a combination of physically-based and deep learning-based protein design methods. Using the Rosetta physically-based protocol from Cao *et al*.^13^, we designed compact (∼60 amino acid) binders for γcommon, βcommon, IFNAR1, and TSLPR, all exhibiting low nanomolar affinities (**Fig. 1A**). For IL12Rβ1, which presents a larger convex binding surface, we applied the modified Rosetta method to produce a 99-residue binder optimized for broad interface engagement^14^. To bind IFNAR2 and IL10Rβ, we used the more recent RFdiffusion method, a deep learning-based approach with a higher success rate in generating functional binders^15^. For IL4Rα and IL2Rβ, we modified the native IL-4 cytokine and the Neo2 IL-2 mimic by computationally mutating their γcommon-binding sites, effectively converting them into single-receptor-binding modules^12^ (**Fig. 1B**). Across the nine receptors, we generated tens of thousands of designs computationally for each and filtered based on AlphaFold’s PAE interaction metric. We screened the subset generated by Rosetta physically-based methods by yeast display due to lower success rates (thousands tested). For designs generated with RFdiffusion we used surface plasmon resonance as we expected higher success rates (96 constructs tested). Across all nine receptors, we generated helical binding proteins, three to five helices each, with binding affinities ranging from 1 to 200 nM (**Fig. 1A, Supplementary Table 1**).

**Figure 1:**
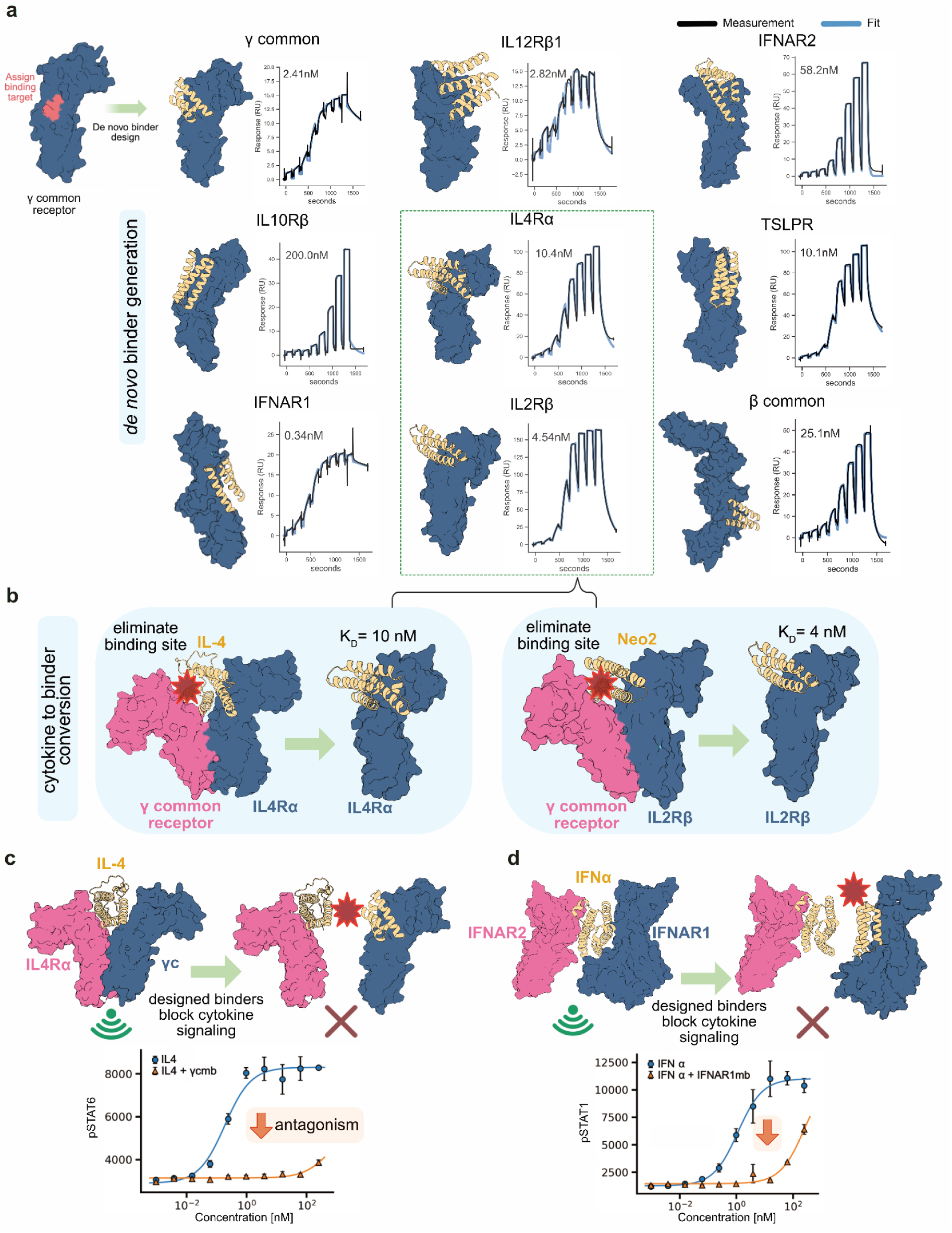
Design and functional analysis of receptor binding proteins. **(a)** Illustration of the nine receptor binding proteins designed in this study, with their binding affinities measured using surface plasmon resonance. Binding affinities are indicated adjacent to each binding curve. **(b)** Overview of the computational method used to transform IL-4 and Neo-2 (IL-2 mimic) agonists into single receptor binders, highlighting the key step of computationally eliminating the γcommon binding site while maintaining the other binding site. **(c)** Illustration of the γcommon binder blocking IL4 signaling by competing for the same binding site (top). 100 nM of IL-4 was added to cells with and without the γcommon binder. Activity was measured after 15 minutes by observing changes in STAT6 phosphorylation via antibody based staining and flow cytometry (bottom). **(d)** Illustration of the IFNAR1 binder blocking strategy (top), Antibody-based STAT1 phosphorylation measurement to assess the IFNAR1 binder capacity to antagonize 100 nM IFNα signaling (bottom).

Since some of these binders target the same interfaces engaged by natural ligands, we hypothesized that they could function as antagonists. When tested on PBMCs, the γcommon and IFNAR1 binders effectively blocked IL-4 and IFNα signaling, respectively **(Fig. 1C-D**). Compared to antibodies, these designed proteins are much smaller, more stable in harsh environments (e.g., the gastrointestinal tract), and amenable to long-term storage^11^. These properties make them promising candidates for therapeutic applications requiring deep tissue penetration (e.g., γcommon in tumors), oral delivery (e.g., IL10Rβ and IFNAR1/2 for gut inflammation), or pulmonary administration (e.g., TSLPR)^16–19^.

### Generating agonists from designed binders

To generate agonists that bring together two receptor subunits, we fused pairs of designed receptor-binding proteins with flexible linkers. In addition to the nine newly designed binders, we incorporated 24 previously developed binders targeting cytokine receptors (IL7Rα, IL10Rα, IL17Rα, IL21Rα, IL23Rα, gp130, IFNLR), TNF receptors (OX40, 4-1BB, TNFR1, TNFR2), immune checkpoints (CTLA-4, PD-L1), and growth factor receptors (cKit, IGF1R, HER2, FGFR, EGFR, BMPR2, ActRII, insulin receptor, TrkA, TGFβR2, ALK1)^10,13,14,18,20–24^. This set of 33 binding proteins allowed us to systematically explore diverse receptor combinations. As an initial test, we reconstructed IL-2 by fusing γcommon and IL2Rβ binders in both fusion orientations while varying linker lengths (**Fig. 2A**). In PBMCs, signaling output depended strongly on orientation (**Fig. 2B**), potentially due to geometric misalignment of intracellular kinase domains. Longer linkers partially alleviated this by adding flexibility, but even the longest (5xGGS linker) could not restore signaling. Solubility was unchanged across variants, supporting geometric alignment as the potential determinant of signaling strength.

**Figure 2:**
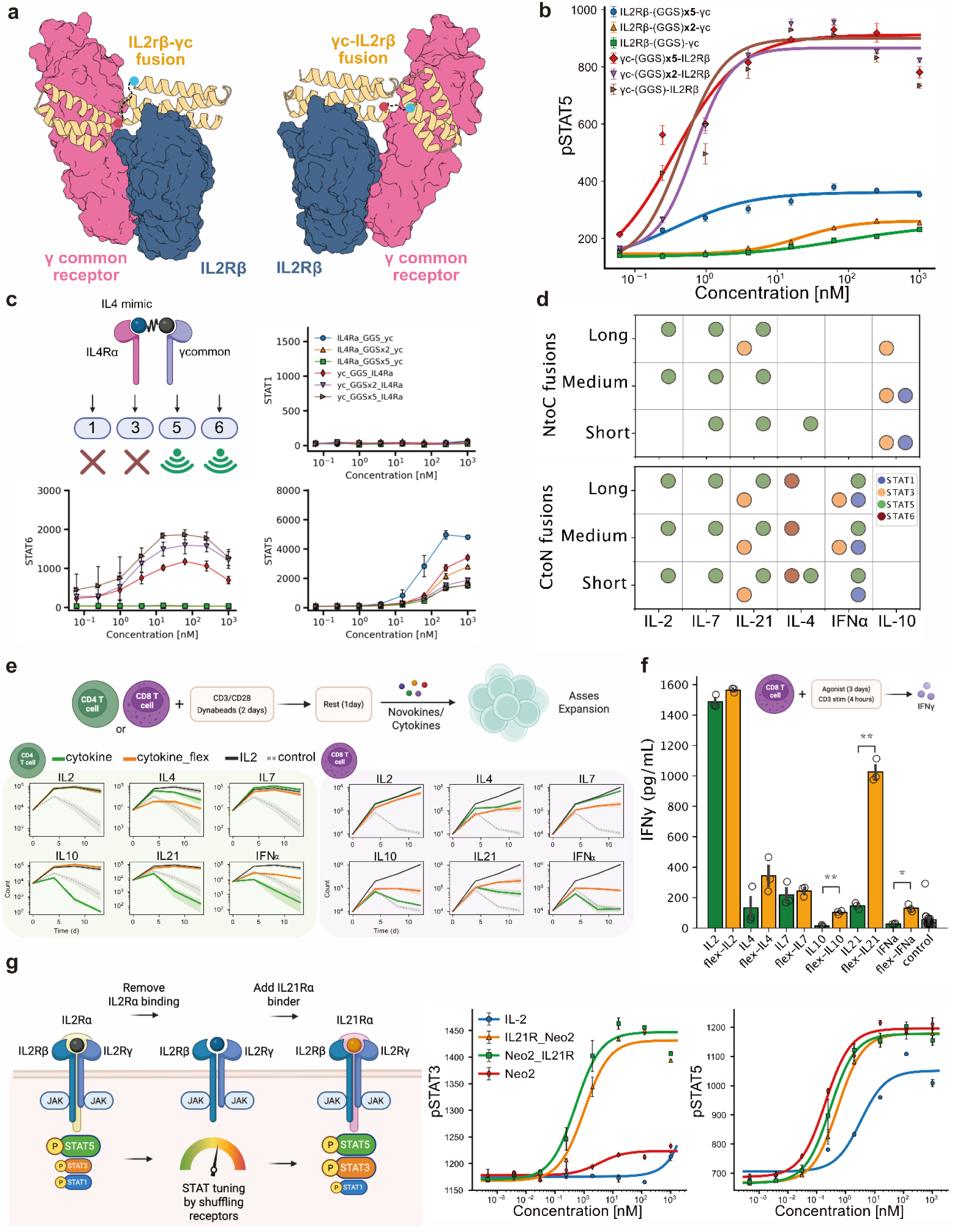
Agonist construction from modular receptor binding domains. **(a)** Schematic of IL-2 analog generation from γcommon and IL2Rβ binders. Two fusion orientations and three linker lengths were tested. **(b)** pSTAT5 signaling in PBMCs following stimulation with the IL-2 mimics for 15 minutes. **(c)** pSTAT1, pSTAT5, and pSTAT6 signaling in PBMCs following stimulation with the IL-4 analogs for 15 minutes. **(d)** Summarized effects of linker length and fusion orientation variations on signaling outcomes of the following mimics: IL-2, IL-4, IL-7, IL-10, IL-21, and IFNα, including pSTAT1, pSTAT3, pSTAT5, and pSTAT6 signaling pathways. **(e)** Assessment of natural cytokines and their novokine analogs to induce proliferation of CD4 and CD8 T cells. Cells were stimulated with CD3/CD28 Dynabeads for two days, rested without beads for one day, and then expanded with the listed cytokine/novokine for each condition. **(f)** Isolated CD8 T cells were incubated with natural cytokines or their analogs for three days and then stimulated with CD3 antibodies for 4 hours to induce IFNy secretion. IFNy secretion was measured with an ELISA. Adjusted *p* values as: ∗: *p* ≤ 0.05; ∗∗: *p* ≤ 0.01. **(g)** Illustration of the strategy used to create an agonist that binds three signaling receptors (left). pSTAT3 and pSTAT5 phosphorylation illustrates that adding an IL21R binding domain to an IL-2 mimic leads to a novel agonist that signals in both STAT pathways (right).

Guided by this principle, we next constructed novokine mimics of IL-4, IL-7, IL-10, IL-21, and IFNα, testing both fusion orientations with medium linkers (2xGGS) for induction of pSTAT1, pSTAT3, pSTAT5, and pSTAT6 (**Fig. 2C-D; Supplementary Fig. 1A**). All synthetic cytokine mimics reproduced the canonical pSTAT pathways of their natural counterparts, confirming that our designs faithfully recapitulate cytokine signaling. However, several mimics went beyond simple reproduction and displayed broadened outputs. For example, an IL-4 mimic activated pSTAT5 in addition to pSTAT6, while IL-21 mimics triggered pSTAT5 alongside pSTAT3 **(Fig. 2D)**. This broadened signaling likely reflects an intrinsic bias toward pSTAT5 when strict dimer geometries are relaxed, given STAT5’s broader pTyr specificity and higher abundance^25,26^. These altered signatures translated into enhanced cellular functions: IL-21 and IL-10 mimics supported CD4^+^ T cell proliferation at levels comparable to IL-2, a capacity absent in their natural counterparts **(Fig. 2E)**. This expanded functionality is most likely linked to the unique STAT5 activation in the mimics. IFNα, IL-10 and IL-21 mimics boosted CD8^+^ T cell proliferation, and IL-21 mimics drove strong IFNγ secretion from CD8+ T cells at levels approaching IL-2 **(Fig. 2E-F)**. Thus, while mimics reproduce native signaling, they can also acquire novel signaling outputs, enabling functions that surpass those of their natural counterparts.

Beyond binary receptor combinations, we extended the strategy to tripartite fusions inspired by IL-2 and IL-15, which naturally bind a third auxiliary receptor without signaling domains. The rationale was that replacing these non-signaling chains with signaling-competent receptors could reprogram downstream outputs. For example, IL-2 primarily drives STAT5, whereas IL-21 drives STAT3. By mixing their receptor components, we could combine or rebalance these transcriptional programs. We generated an IL-2/IL-21 hybrid by incorporating an IL-21Rα binding domain into Neo2, a de novo IL-2 analogue (**Fig. 2G**). This chimera activated both pSTAT5 and pSTAT3, unlike IL-2 or Neo2, which mainly activated pSTAT5. Because pSTAT3 promotes T cell survival, stem-like memory, and resistance to exhaustion, this hybrid could enable new functionalities absent in the IL-2 cytokine^27,28^. We refer to this class as Multikines: modular cytokines designed to engage multiple receptors. Multikines provide a generalizable strategy for constructing novokine libraries that systematically map and engineer signaling landscapes beyond dual-receptor pairings.

### Large scale agonist generation and characterization

With a modular set of receptor-binding domains and a fusion strategy, we set out to map the signaling landscape of novokines. In an initial pilot of 576 fusions from 24 binders expressed in a high-throughput bacterial system, we detected specific, non-promiscuous activation in pSTAT1/3/5/6 pathways. These patterns recapitulated signaling by IL-2, IL-4, IL-7, IL-10, IL-21, and IFNα, validating both the sensitivity and specificity of our approach **(Supplementary Fig. 2A, 3)**. Encouraged by this, we expanded to a larger library of 1,089 novokines spanning all 33 receptor binders and tested them across two donors **(Fig. 3A; Supplementary Fig. 2B-C)**. For one donor, we extended the staining panel to thirteen pathways **(Supplementary Fig. 2B)**. Single binder controls were inactive, confirming the requirement for dual-receptor engagement. From the full set, 75 fusions triggered reproducible pSTAT responses in at least two out of three donors. Constructs involving γcommon exhibited the highest activation rate (42%), but other receptors, including gp130, IL7Rα, IL10Rα, IL21Rα, βcommon, TrkA, and IFNAR1, also signaled in ∼20% of cases. Several of these are not traditionally considered shared components, highlighting untapped signaling potential.

**Figure 3:**
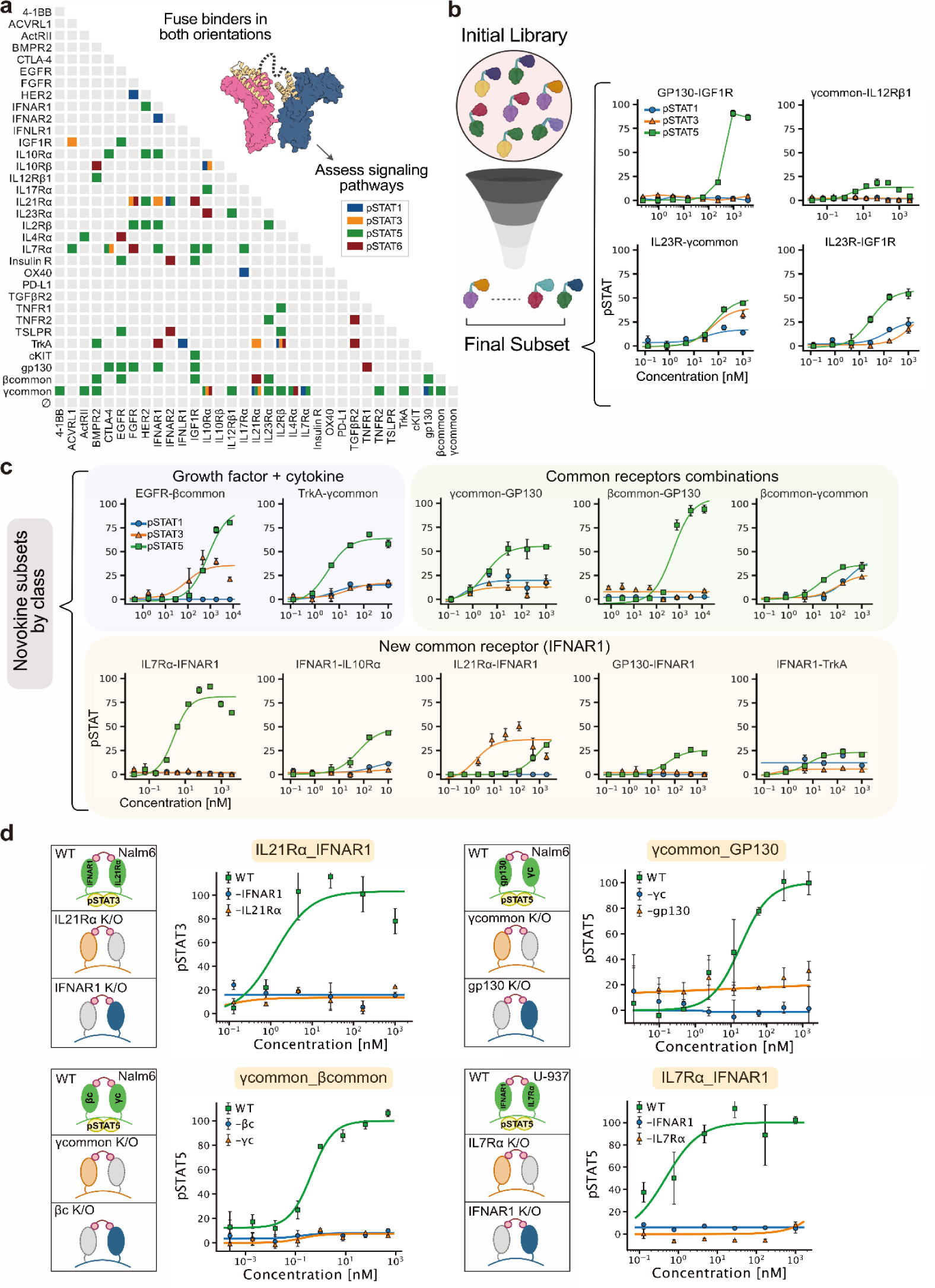
Large scale evaluation of STAT phosphorylation by a library of designed agonists. (**a**) Signaling activity of novokines generated by all-by-all fusions of receptor binders. Only ligands exceeding two standard deviations above the control mean are shown. (**b**) Dose-response validation of fourteen novokines from (**a**) across pSTAT1, pSTAT3, and pSTAT5 pathways. (**c**) Representative agonists grouped into three categories: (1) cytokine-growth factor fusions, (2) shared subunit fusions, and (3) IFNAR1-based combinations. (**d**) Receptor dependence confirmed by CRISPR-Cas9 knockout of individual subunits. STAT phosphorylation responses were lost when either receptor partner was deleted, demonstrating that signaling requires the designed receptor pair. Knockouts were validated by absence of responses to the corresponding natural cytokine.

As with any high-throughput screen, potential artifacts required careful evaluation. Signaling could in principle arise from endotoxin contamination or from clustering driven by self-association of a single binding domain. To rule out endotoxin effects, we tested high levels of endotoxin (up to 1000 ng/ml, far exceeding sample content produced under low-endotoxin conditions using CHAPS detergent) and observed no pSTAT activation during short 15-minute incubations **(Supplementary Fig. 3)**. To assess clustering, we performed dose-response experiments on 20 constructs. Fourteen showed saturable, dose-dependent signaling consistent with a dual-receptor mechanism, four displayed non-saturating responses consistent with clustering, and three failed to signal strongly enough to clearly see a dose-dependent response **(Fig. 3B-C; Supplementary Fig. 4A)**. To test the dependence on the targeted receptor subunits, we selected four novokines (βcommon-γcommon, γcommon-gp130, IL7Rα-IFNAR1, IL21Rα-IFNAR1), and knocked out each of the corresponding receptor subunits. For each of the four, knockout of either receptor subunit eliminated signaling, demonstrating that signaling requires both targeted receptor subunits (**Fig. 3D)**. Together, these controls show that most tested novokines act through their designed receptor pairs, as evidenced by saturating dose curves and confirmed by knockouts in four cases.

From this screen, we identified fourteen novokines with saturating dose-response curves that fall into three mechanistic classes. The first comprised fusions of common receptors with one another, such as βcommon-γcommon, underscoring the central role of shared subunits in productive cytokine signaling; notably, no pairs between private cytokine receptors were detected. The second linked cytokine receptors with non-cytokine partners, including growth factor receptors such as TrkA, an unusual but not unprecedented architecture that highlights underappreciated cross-family compatibility^29,30^. While both EGFR and TrkA are expressed at relatively low levels in T cells, macrophages, and other immune subsets, multiple studies show that they remain functionally active and capable of modulating cytokine production, activation, and inflammatory responses^31–38^. The third class involved private cytokine receptors paired with IFNAR1. Although IFNAR1 normally dimerizes only with IFNAR2, these constructs revealed an unexpected ability of IFNAR1 to act as a versatile signaling scaffold, resembling a common receptor. We focused on this third IFNAR1-based subset as a new axis for expanding cytokine signaling architectures, as described in the following section.

### IFNAR1 novokines

Beyond its native partner IFNAR2, IFNAR1 supported signaling with multiple receptor chains, including IL-21Rα, IL-7Rα, IL-10Rα, and gp130 **(Fig. 4A)**. This broad pairing capability suggested that IFNAR1 may function analogously to established shared receptor subunits such as γcommon and βcommon. As with other common receptors, in all signaling novokines IFNAR1 served as the subunit with the shorter intracellular domain (a feature of common receptors; **Supplementary Fig. 5**), and its relatively low affinity for type I interferons is consistent with the paradigm of low-affinity common receptors pairing with high-affinity private ones^39^.

**Figure 4:**
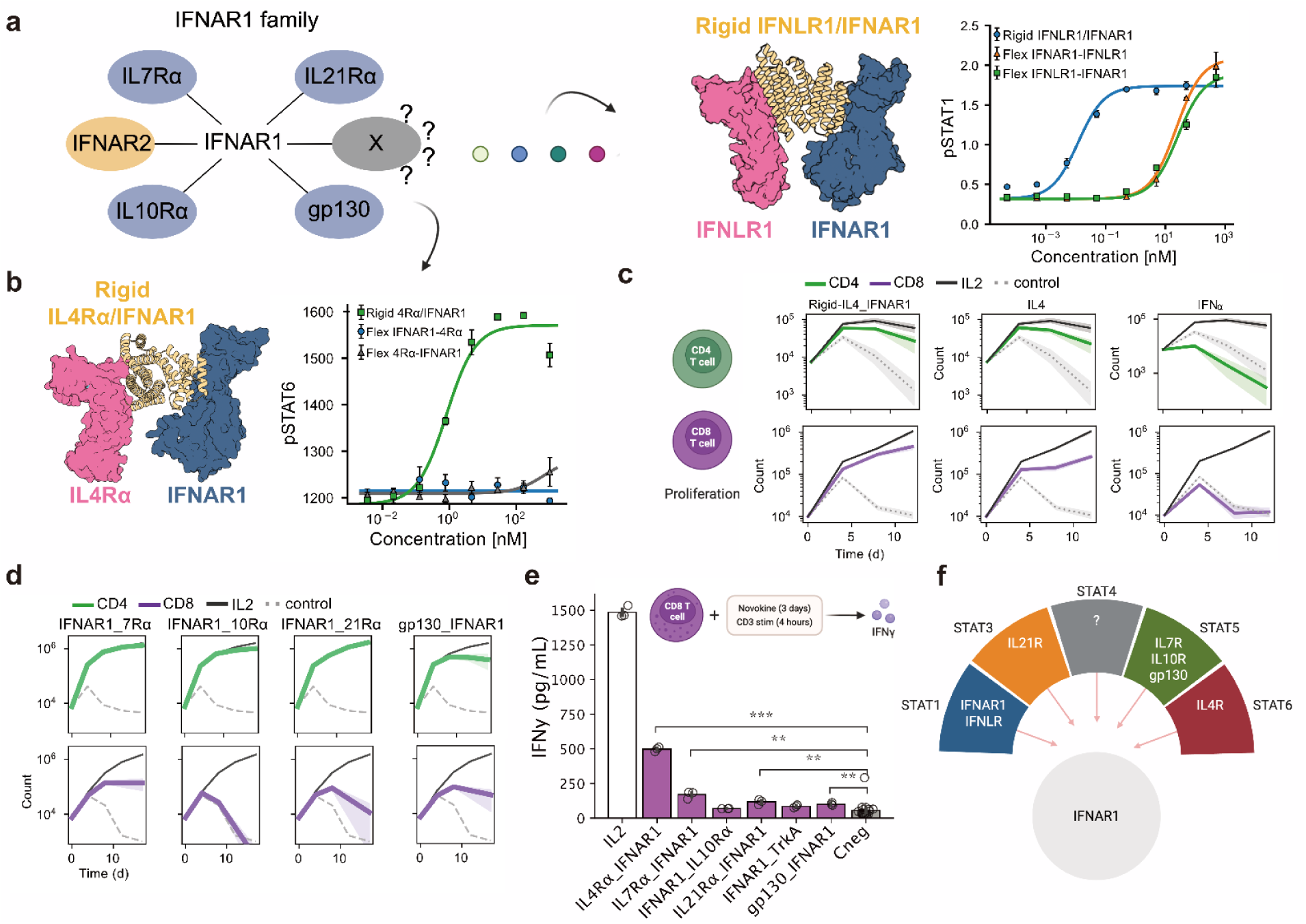
Characterization of a novel IFNAR1 synthetic signaling family. **(a)** Identification of IFNAR1 as a previously unexplored common receptor in our screen opened up the possibility that it could function as a shared receptor (left). When paired with IFNLR, IFNAR1 was able to replace the IL10Rβ common receptor and recapitulate signaling in STAT1 (right). Rigidification of the ligand was needed to enhance signaling activity. **(b)** When paired with IL4Rα, IFNAR1 was able to substitute for the γcommon receptor and restore pSTAT6 signaling. Ligand rigidification was required to improve potency. **(c)** Assessment of the IL4/IFNAR1 novokine for its capacity to induce proliferation of CD4^+^ and CD8^+^ T cells along with the IL4 and IFNα cytokines**. (d)** Assessment of selected novokines from the IFNAR1 family for their capacity to induce proliferation of CD4^+^ and CD8^+^ T cells. **(e)** CD8^+^ T cells were incubated with the IFNAR1 family of novel ligands for three days and later stimulated with CD3 antibodies for 4 hours to induce IFNy secretion. IFNy secretion was measured via ELISA. Adjusted *p* values as: ns: *p* > 0.05; ∗: *p* ≤ 0.05; ∗∗: *p* ≤ 0.01; ∗∗∗: *p* ≤ 0.001. (**(f)** The novel family generated around IFNAR1 can signal in multiple STATs based on the receptor chosen to dimerize, leading to a variety of potential functional effects.

To probe this modularity further, we tested IFNAR1 in a combination with IFNLR, a receptor which is absent in PBMCs. In HEK cells overexpressing IFNLR, IFNAR1-IFNLR fusions triggered pSTAT1 activity, but only at high ligand concentrations. Building on our companion work where rigidly constraining receptor geometry enhanced signaling, we applied de novo design to enforce a rigid IFNLR-IFNAR1 geometries that restored potent pSTAT1 activity^40^**(Fig. 4A)**. Using the same strategy, we recovered pSTAT6 signaling from IL-4Rα-IFNAR1, showing that enforced geometry can unlock outputs inaccessible to flexible fusions and expand the IFNAR1 signaling repertoire **(Fig. 4B)**.

To assess the functional consequences of IFNAR1-based novokines, we tested their effects on T cell proliferation. IL-4Rα-IFNAR1 drove both CD4⁺ and CD8⁺ T cell proliferation at levels closely resembling IL-4, diverging from the limited effects of IFNα **(Fig. 4C)**. This indicated that the private receptor IL-4Rα shaped the functional outcome, consistent with it shaping the pSTAT6 signature. Other IFNAR1 pairings (IL-7Rα-IFNAR1, IL-21Rα-IFNAR1, IL-10Rα-IFNAR1, gp130-IFNAR1) also supported CD4⁺ T cell proliferation, with IL-7Rα-IFNAR1 extending partially to CD8⁺ T cells **(Fig. 4D)**. Beyond proliferation, IL-4Rα-IFNAR1 elicited the strongest IFNγ secretion in activated CD8⁺ T cells among all IFNAR1 fusions **(Fig. 4E)**. This functional breadth suggests that IFNAR1 can act as a synthetic signaling hub, analogous to γcommon, with broad pSTAT signaling signatures and potential for immune modulation **(Fig. 4F)**.

### Common receptors are signaling hubs and private receptors shape signaling

When pairing IL-4Rα with IFNAR1 we generate a signaling and functional output that mirrors IL-4 suggesting a dominant role for the private receptor while highlighting flexibility in the choice of common receptor. To further probe these observations, we next asked whether IL-4Rα could also engage additional common partners, and if so, whether it would continue to dictate the signaling outcome. To test this, we rigidly paired IL-4Rα with IL-10Rβ and βcommon, which had failed to pass our positive signal thresholds in the flexible screen. Both constructs successfully induced pSTAT6 phosphorylation **(Fig. 5A-B)**, supporting two principles: common receptors are modular and swappable across cytokine families, and signaling identity is majorly dictated by the private receptor **(Fig. 5C)**.

**Figure 5:**
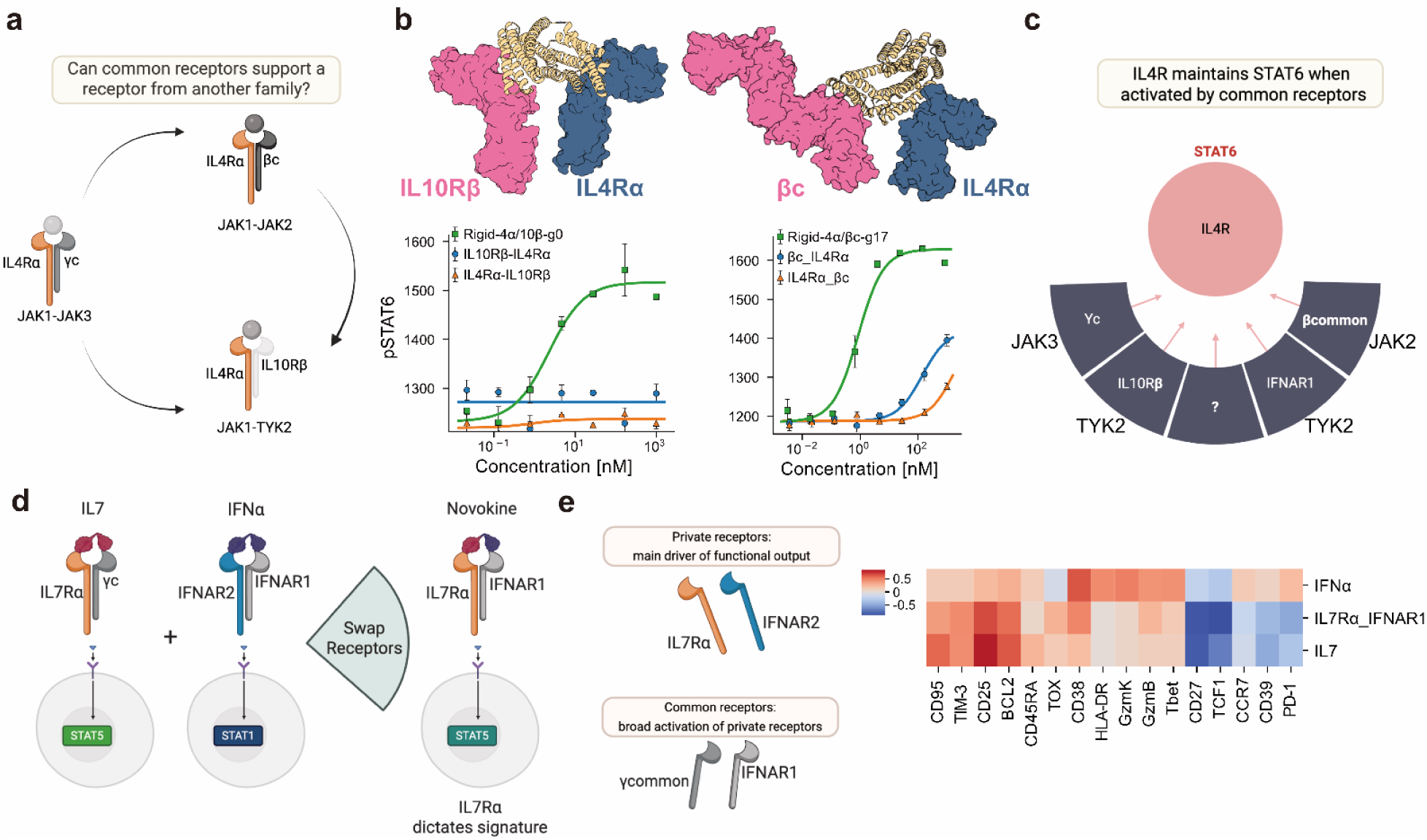
Common receptors are signaling hubs and private receptors shape signaling. **(a)** The ability of IL4Rα to signal with IFNAR1 presents a hypothesis where other common receptors can be dimerized with IL4Rα to generate a pSTAT6 signal. **(b)** Rigidified novokines dimerizing IL4Rα with IL10Rβ and βcommon receptors respectively induced STAT6 signaling. **(c)** IL4R can serve as a STAT6 signaling node regardless of which common receptor is paired with it. (**d**) Illustration of a synthetic IFNAR1-IL7Rα novokine created by combining binders for IL7Rα with IFNAR1. **(e)** Expression profiles of cell markers in naive CD8^+^ T cells activated with CD3/CD28 Dynabeads shows that the private receptor IL7Rα, like IL4Rα, shapes not only the canonical STAT signature but also broader transcriptional programs, demonstrating that private receptors are the key determinant of signaling outcomes and common receptors serve as activators but minimally shape the signal.

Finally, to test whether private receptor dominance extends beyond IL-4Rα, we examined the IL-7Rα-IFNAR1 novokine, which signals through pSTAT5 analogous to IL-7, and assessed its ability to shape protein expression in activated CD8⁺ T cells. The resulting profile, assessed by flow cytometry, closely resembled that of IL-7, with some variations, and diverged from the profile induced by IFNα **(Fig. 5D-E)**. Altogether, these findings support a unifying principle: private receptors define signaling and functional identity, while common receptors provide flexible assembly platforms that expand cytokine architecture without fundamentally altering the response.

### Functional characterization of Novokines

While these findings establish that private receptors define the signaling identity of a novokine, the extent to which novokines can enable cell type-specific responses remains unknown. To address this, we profiled novokine activity across PBMCs from three donors and observed that novokines frequently diverged from both their natural cytokine counterparts and from one another in the immune subsets they engaged, forming three different behaviors **(Fig. 6A; Supplementary Fig. 6)**. First, several novokines gained new activities. IL-7Rα-IFNAR1 extended IL-7 signaling beyond T cells to also drive pSTAT5 in NK cells, gp130-IFNAR1 also promoted similar T and NK cell responses. IL-21Rα-IFNAR1 activated pSTAT3 broadly across T cells, B cells, and dendritic cells (DCs). Second, some novokines lost or shifted specificity. IL-21 and IL-10 novokine mimics lost pSTAT3 signaling in DCs and B cells, respectively, while IL-4Rα-IFNAR1 restricted pSTAT6 activity to B cells and DCs mainly compared to the broader signaling of IL-4. Third, certain pairings produced unexpected outputs. IL-4Rα-IL-10Rβ and IL-4Rα-βcommon were similar in pSTAT6 activation overall, except that IL-10Rβ pairing uniquely induced pSTAT6 in monocytes, whereas βcommon pairing redirected signaling toward pSTAT5 in T and NK cells. Novokines incorporating receptors with low abundance in PBMCs (IL-12Rβ1, TrkA) produced pSTAT5 activity that was largely confined to naïve and central memory T cells **(Supplementary Fig. 7)**. Given the low expression of these receptors in PBMCs, it remains possible that the observed activity arises through mechanisms such as receptor superclustering, which will require further investigation. Together, these findings show that novokines can generate cell-type-specific signaling and that the same novokine can produce distinct signatures depending on the responding population.

**Figure 6:**
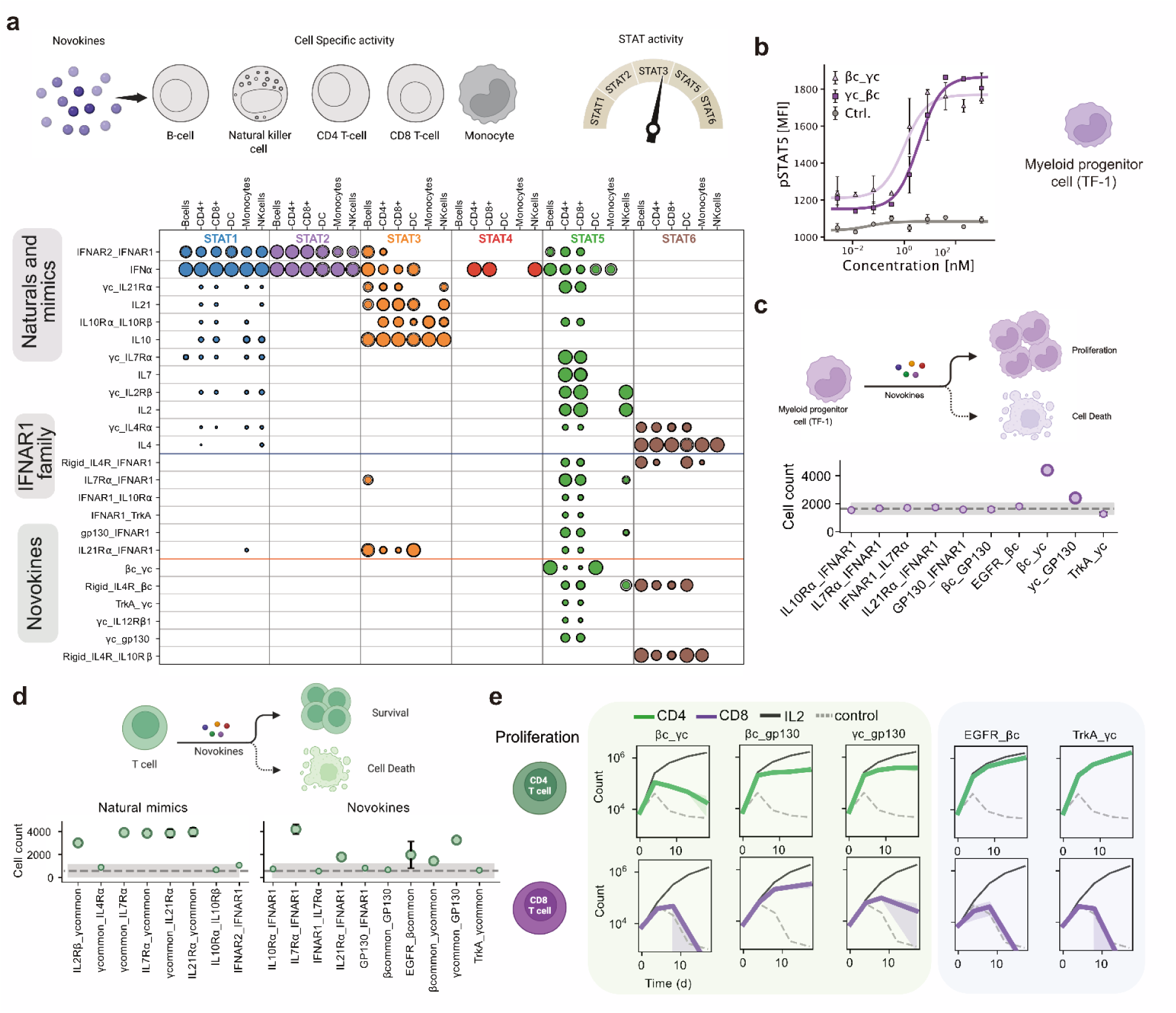
Characterization of novokine signaling. (**a**) Cell-type-specific activity of selected novokines in primary PBMCs across pSTAT1, pSTAT3, pSTAT4, pSTAT5, and pSTAT6 pathways, tested independently in three donors. **(b)** The βcommon-γcommon ligand showed weak activity in bulk PBMCs and was further tested in a monocyte cell line (TF-1). **(c)** Evaluation of novokines for their ability to drive TF-1 proliferation. Cells were expanded in the presence of the novokines for three days. **(d)** Evaluation of novel agonists for their ability to support naive T cell survival. Cell counts were read with flow cytometry eight days after assay initiation. **(e)** Assessment of selected novokines from two classes for their capacity to induce proliferation of CD4 and CD8 T cells.

We next asked whether these signaling differences translated into lineage-specific functions. We focused on the TF-1 cell line, a model of the myeloid lineage, which relies on βcommon signaling for proliferation. To probe if novokines can support this function, we first tested βcommon-γcommon in TF-1 cells, and observed robust, dose-dependent pSTAT5 activation. **(Fig. 6B)**. We then probed whether it could substitute for IL-3 or GM-CSF, both of which are used to activate βcommon for TF-1 proliferation. Under IL-3 depleted conditions, βcommon-γcommon was the most effective novokine amongst the tested subset at sustaining cell growth **(Fig. 6C)**, establishing its functional capacity for this lineage.

Having established that βcommon-γcommon can support proliferation in a myeloid cell line, we next turned to naïve T cells. Because IL-7 is central to T cell homeostasis, we asked whether the IL-7Rα-IFNAR1 novokine could similarly support survival and proliferation. Naïve T cells were cultured with novokine mimics of natural cytokines as well as select novel pairings, including IL-7Rα-IFNAR1. Among natural mimics, survival was promoted by IL-2, IL-7, and IL-21 mimics **(Fig. 6D)**. The IL-7Rα-IFNAR1 novokine also supported survival and produced the strongest response, followed by γcommon-gp130. To evaluate functional effects in broader T cell subsets, we stimulated CD4⁺ and CD8⁺ T cells with CD3/CD28 and expanded them with ligands from the shared-subunit fusions and growth factor-cytokine hybrids. βcommon-γcommon did not expand T cells, consistent with its weak T cell activity. EGFR-βcommon, TrkA-γcommon, and γcommon-gp130 on the other hand, selectively expanded CD4⁺ but not CD8⁺ T cells, while βcommon-gp130 promoted proliferation of both subsets **(Fig. 6E)**.

Together, these results demonstrate that novokines can target specific immune populations and support specialized functions: βcommon-γcommon supported TF-1 cell proliferation, IL-7Rα-IFNAR1 promoted naïve T cell survival, and several novokines such as TrkA-γcommon preferentially expanded CD4⁺ versus CD8⁺ subsets.

## Discussion

By assembling 33 modular receptor-binding domains in an all-by-all fashion, we generated and tested more than 1,000 de novo designed agonists spanning over 500 unique receptor pairings. This effort uncovered more than eighteen previously unreported signaling pairs in primary immune cells and revealed three broad mechanistic classes: (i) shared subunit fusions such as γcommon-gp130, (ii) cytokine-growth factor hybrids such as TrkA-γcommon, and (iii) an IFNAR1-based axis that displayed remarkable versatility across diverse partners. The designed binders are structurally programmable, allow control over orientation and linker geometry, and can be produced at high throughput in bacteria, making them ideally suited for systematic exploration of receptor space.

The scale of our screen, the diversity of receptor pairings tested, and the use of primary human immune cells together represent a substantial advance over previous efforts to engineer novel cytokine signaling. Earlier studies showed that non-natural receptor dimers could trigger signaling in engineered cell lines, but these efforts were limited in scope and did not reflect native cellular contexts^4^. More recently, synthetic cytokines were shown to induce new biological functions in engineered primary cells, highlighting the potential of this concept, but that approach required genetic modification of immune cells to overexpress synthetic receptors^8^. Similarly, nanobody-based bispecific cytokines have been used to dimerize receptor pairs in primary cells, but nanobodies depend on immunization and large-scale screening, offer limited control over geometry, and are not easily extended to systematic mapping^5,41^. In contrast, our approach combines de novo protein design with high-throughput functional screening and controllable geometrical design, enabling rapid, programmable exploration of a broad receptor space at a scale unattainable with antibody-based formats.

Several general principles emerged from our screen. First, pSTAT5 was the most readily activated pathway across PBMCs, often triggered by diverse receptor combinations, consistent with its abundance and broad substrate specificity. Second, established shared subunits such as γcommon, βc, and gp130 were repeatedly co-opted in non-natural pairings, underscoring their modularity. Third, IFNAR1 emerged as an unexpected common receptor, able to pair with IL4Rα, IL7Rα, IL21Rα, IL10Rα, IFNLR, and gp130 to activate distinct STAT programs (pSTAT1, pSTAT3, pSTAT5, and pSTAT6). Fourth, we identified receptor fusions that recruited JAK/TYK combinations not found in nature, such as JAK3/JAK2 with βcommon-γcommon and TrkA-γcommon linking cytokine and growth factor pathways. These findings suggest that the natural cytokine signaling network may reflect evolutionary constraints rather than absolute biochemical limits, pointing to new opportunities for immune modulation.

Novokines produced distinct cell type-specific responses: βcommon-γcommon preferentially activated B cells, monocyte-like cell lines, and DCs, whereas IL7Rα-IFNAR1 promoted T cell proliferation. Overall, novokine signaling was weaker than that of native cytokines, which we attribute to partial or unstable JAK engagement. However, this limitation was overcome by rigidly constraining receptor geometry, which both strengthened signaling and restored activity where flexible designs failed. Furthermore, whereas natural cytokines are optimized to act within receptor families, novokines expand what is possible by creating cross-family pairings that begin to define a synthetic signaling landscape enabling cell-specific and tunable immune responses **(Fig. 7)**.

**Figure 7:**
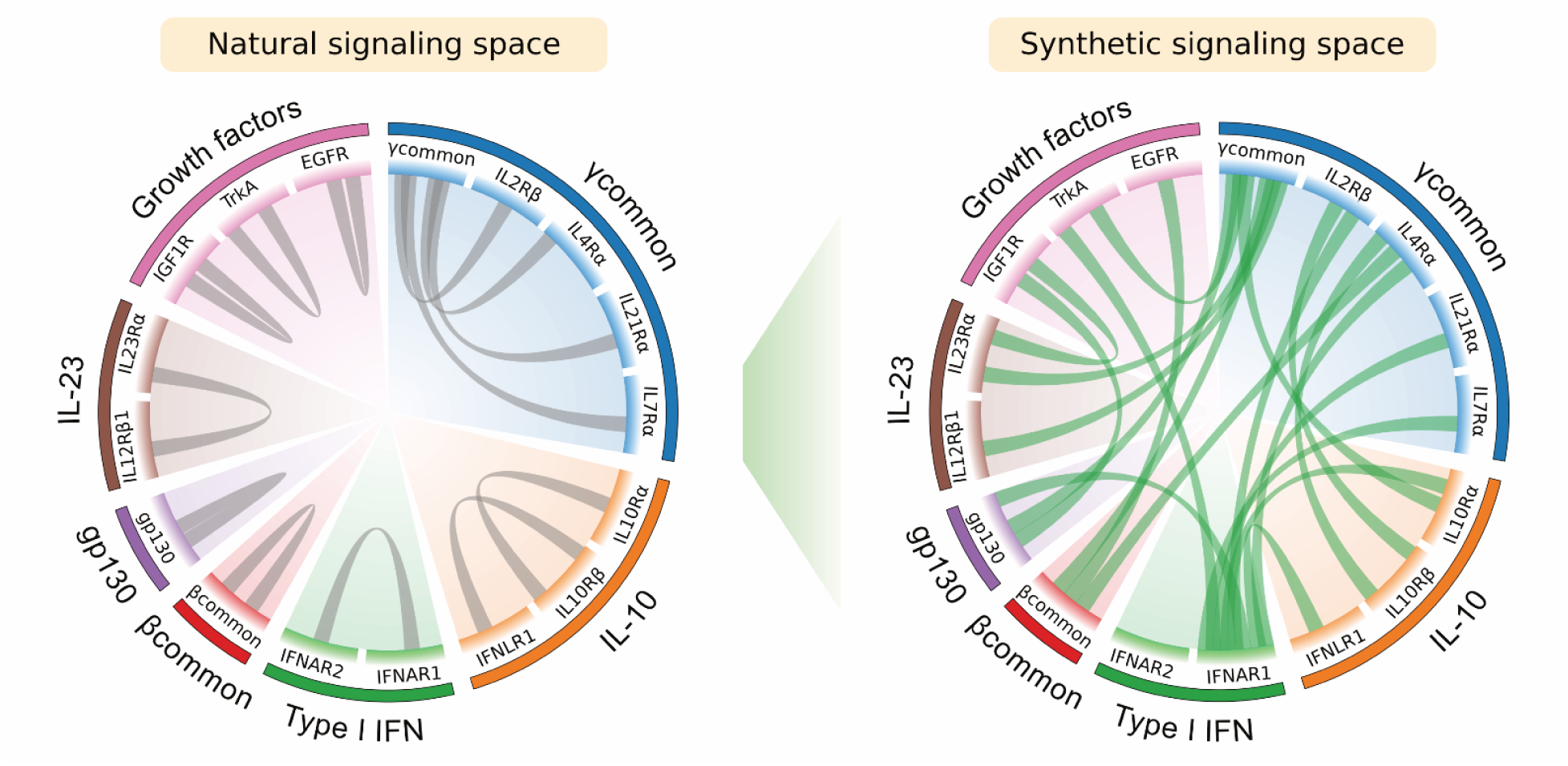
Expanding the cell signaling space with Novokines. Illustration of the natural signaling pairs accessible by a subset of the receptors tested that successfully generated Novokines (left).The expanded synthetic signaling space of functional pairs verified through dose curves (right).

Our results demonstrate that receptor pairings determine both the cell type specificity and the signaling pathways accessible to a cytokine complex. Within this framework, private receptors dictate signaling identity, while common receptors serve as modular scaffolds that enable diverse pairings. Consistent with this, most productive novokines require at least one common receptor, though the molecular features that make common receptors uniquely suited for this role remain to be defined. Elucidating these features may uncover new principles of cytokine biology. From a design standpoint, systematically sampling receptor combinations provides a strategy to generate cell type-specific signaling signatures, which can then be further refined through geometric control. Together, these findings through our screen reveal a more flexible and designable cytokine architecture than exists in nature, with broad potential for therapeutic immune modulation.

## Limitations of the study

As with any high-throughput effort, our agonist screen has several limitations. First, variability across human donors may influence outcomes. While we focused on responses reproducible across three donors, broader cohorts could reveal additional heterogeneity, particularly in rare subpopulations that may be underrepresented in bulk PBMC assays. Second, although our designed binding domains exhibit reasonable affinity and specificity, off-target interactions with other surface proteins cannot be fully excluded. For four novokines (IFNAR1-IL7Rα, IFNAR1-IL21Rα, βcommon-γcommon, and gp130-γcommon), receptor knockouts confirmed signaling through the intended receptor pairs, but more comprehensive studies will be needed to confirm the signaling mechanism. Other hits from this screen should undergo further mechanistic analysis to better map and understand the synthetic signaling space. In particular, novokines signaling through receptors with low expression in PBMCs, such as TrkA or EGFR, will require additional work to clarify whether their signaling outputs reflect unintended mechanisms such as receptor clustering or other context-dependent effects. Third, our signaling analysis was restricted to a single time point snapshot of a few pathways. Because cytokine responses vary in timing and magnitude, dynamic time-course experiments or transcriptome-wide profiling will be necessary to capture the full range of activities. Finally, resting PBMCs in vitro do not recapitulate the complexity of physiological environments such as inflamed tissues or the tumor microenvironment. Future studies that test novokines across diverse cellular states and contexts will be essential to fully understand their functional potential.

## Acknowledgements

We thank members of the Baker lab for helpful discussions and feedback on the project, L. Goldschmidt and P. Vecchiato for computational infrastructure, and K. VanWormer and H. Nunez-Ortega for wet-lab infrastructure. We are grateful to J. Decarreau, S. Cheng, C. Dobbins, and members of the bioassay core for preparing PBMCs and assistance with cell lines, and B. Wicky, L. Milles, R. Ragotte, J. Qian, and S. Gerben for technical support with protein expression and purification. Protein visualization was done in PyMol. Some schematic figures included in this work were created with or modified from BioRender. We gratefully acknowledge the support of numerous organizations and funding agencies that made this work possible. This research was supported by the Howard Hughes Medical Institute (D.B.), the National Institutes of Health (NIH), including the National Cancer Institute (R01CA114536 and R01CA240339); and the Bill & Melinda Gates Foundation (INV-043758). Additional support was provided by the IARPA Genetic Circuits program, and the DOD Breakthrough Award W81XWH-20-1-0230. This work further benefited from the Open Philanthropy Project Improving Protein Design Fund, the Audacious Project, the Nordstrom Barrier Institute for Protein Design Directors Fund, the IPD Design Fund, and the Washington Research Foundation Protein Design Hub. M.H.A. is a Fellow of The Jane Coffin Childs Fund for Medical Research. M.E. acknowledges support from the “La Caixa” Foundation (ID 100010434 under grant no. LCF/BQ/AA19/11720031), and the “Rafael del Pino” Foundation.

## Author Contributions

M.H.A. and M.E. conceived and designed the initial project, interpreted data, and wrote the manuscript. M.H.A., M.E, B. C., A.K., J.C., M.L., T.Y. M.G., and I.G. designed binders. D. M. ran the phenotyping experiments in Figure 6A and analyzed the data. S.J. performed experiments and interpreted data. D.B. supervised experiments and reviewed and edited the manuscript.

## Data Availability

All raw data and analysis associated with this study not currently present within the manuscript and SI figures will be provided within supplementary information files.

## Conflict of interest

D.B., M.H.A., M.E., B.C, A.K., T.Y., M.G., M.L.-H., and I.G. are inventors of a provisional patent application submitted by the University of Washington for the designed novokines. E.J.W. is a member of the Parker Institute for Cancer Immunotherapy which supported this study. E.J.W. is an advisor for Arpelos Biosciences, Arsenal Biosciences, Coherus, Danger Bio, IpiNovyx, New Limit, Marengo, Pluto Immunotherapeutics, Related Sciences, Santa Ana Bio, and Synthekine. E.J.W. is a founder of Arpelos Biosciences, Danger Bio, and Arsenal Biosciences. E.J.W. holds stock in Coherus.

## Supplementary Figures and Tables

**Supplementary Figure 1:**
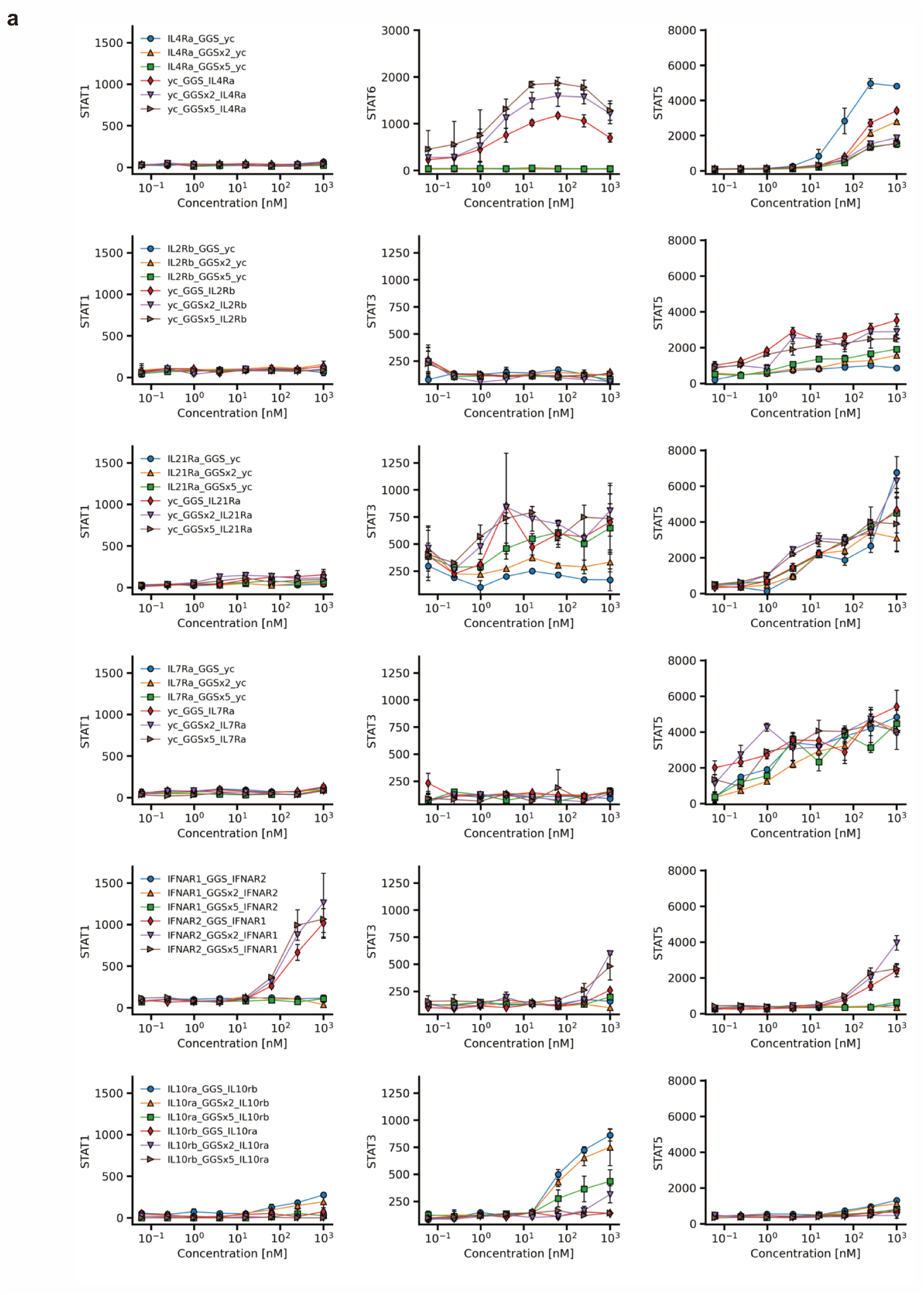
pSTAT signaling of novokine mimics of natural cytokines. (a) dose-response curves for pSTAT1, pSTAT3, pSTAT5, and pSTAT6 signaling that was used to generate the grid plot shown in Figure 2D. Ligands were tested in PBMCs that were stimulated for 15 minutes.

**Supplementary Figure 2:**
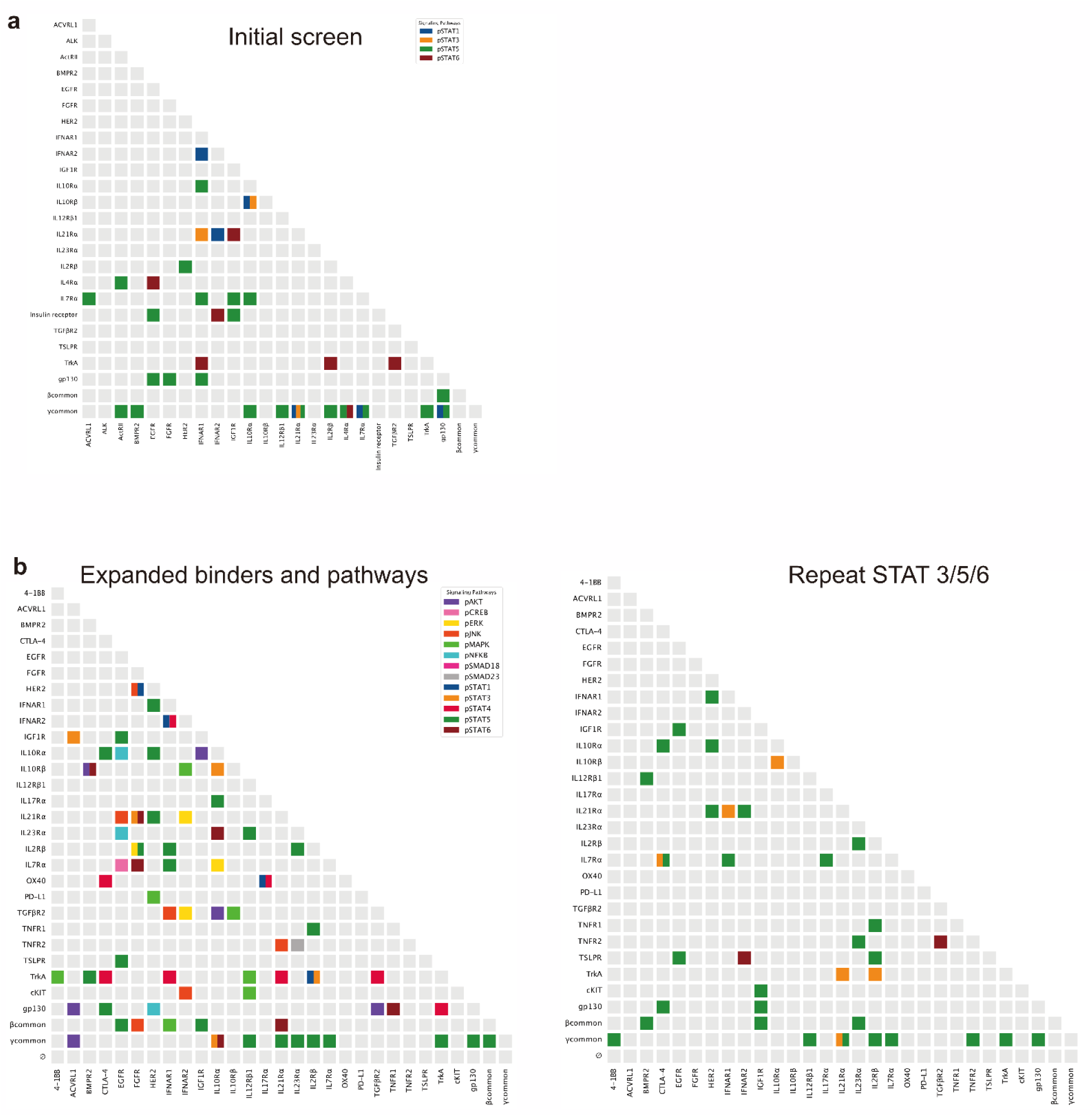
All-by-all signaling activity from three experiments. (a) Initial results from signaling screen across the pSTAT1,3,5,6 pathways for the initial pilot of 576 fusions. (b) Results from signaling screen across 13 different signaling pathways and the full library of fusions. IL4R and Insulin receptor proteins are missing since they failed to transform in this run. (c) Repeating the experiment from (b) in an independent PBMC donor while assessing the pSTAT3,5,6 signaling pathways. pSTAT1 was attempted but the antibody batch used failed to capture positive controls and as a result was omitted.

**Supplementary Figure 3:**
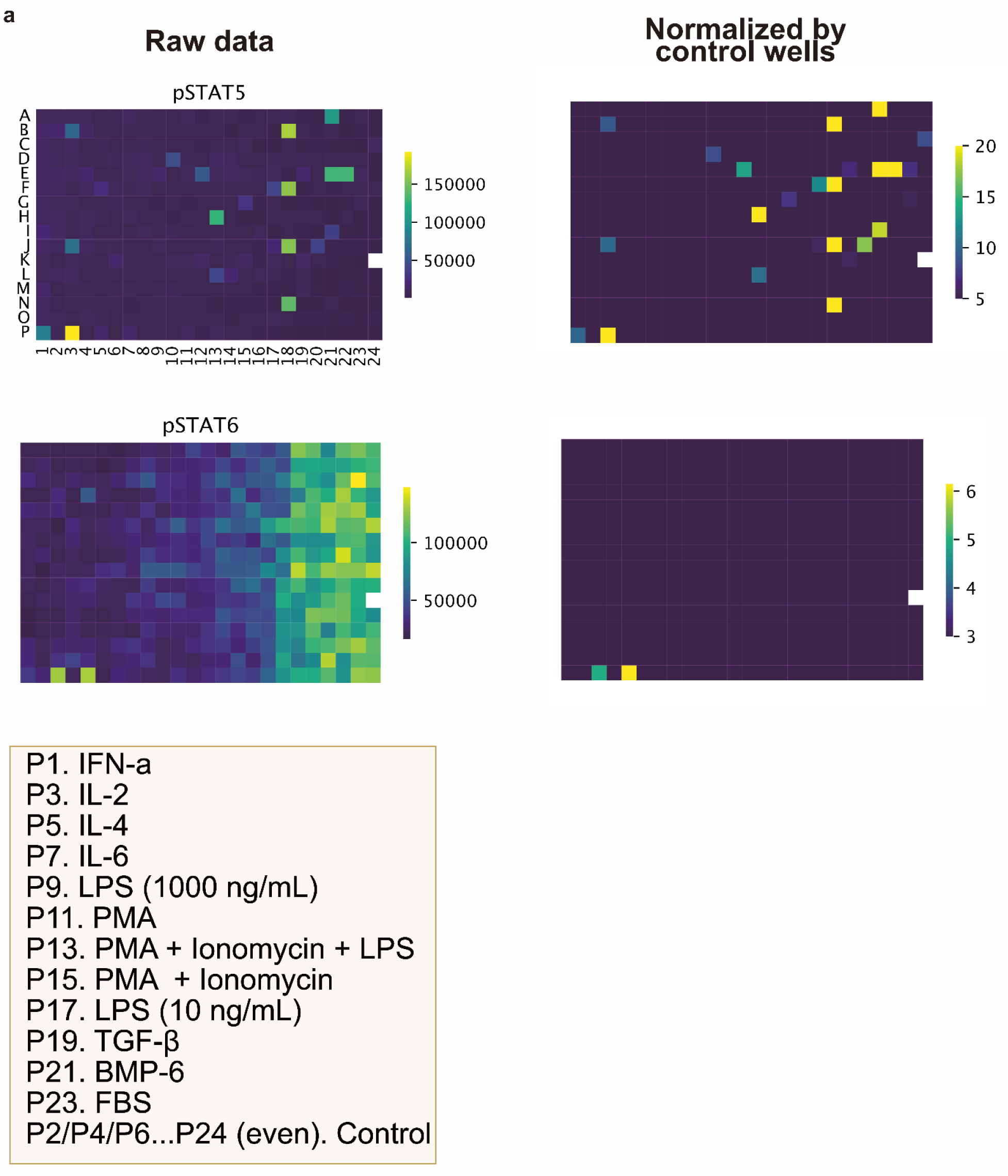
Example of an all-by-all signaling activity plate measurement. (a) Signaling results from the all-by-all analysis across the STAT5,3,6 pathways for a subset of the data shown in Figure 3A. To control for batch effects, we included negative controls across each plate and pathway-specific positive controls to benchmark signaling responses.

**Supplementary Figure 4:**
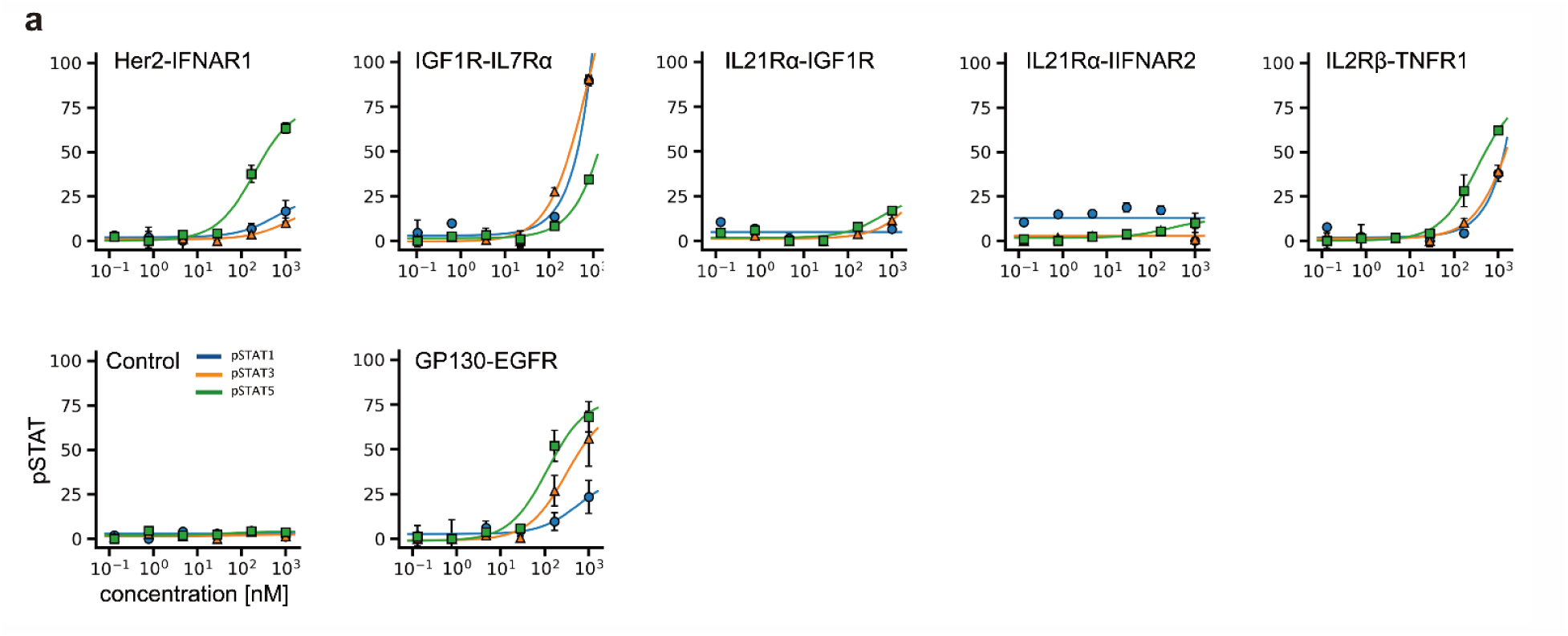
Novokines that failed to pass further verifications. (a) Signaling results from the all-by-all analysis across the pSTAT1,3,5, 6 pathways for a subset of the seven hits that failed verifications test due to non saturating signals, or lack of signaling.

**Supplementary Figure 5:**
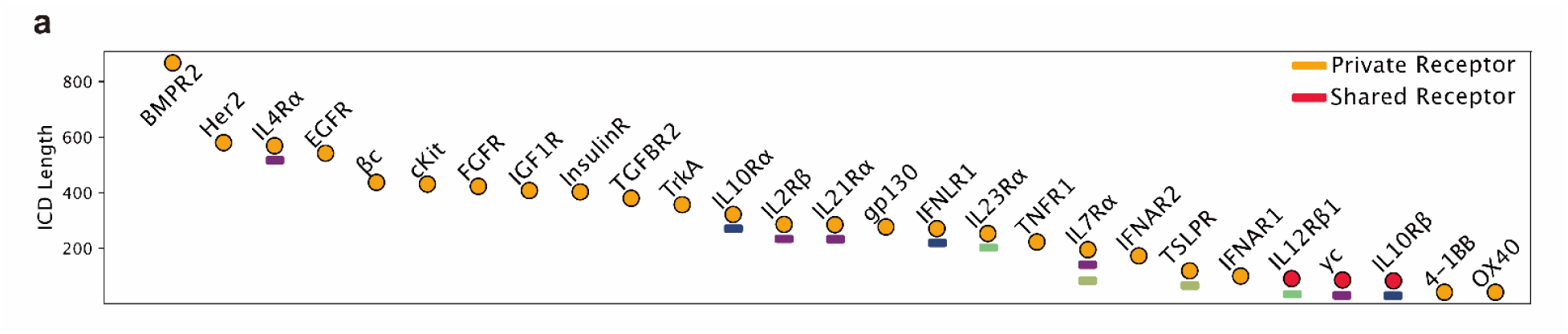
ICD length across select receptors. **(a)** common receptors that form heteromeric signaling complexes are marked with a red circle. Lines under receptors match known pairs.

**Supplementary Figure 6:**
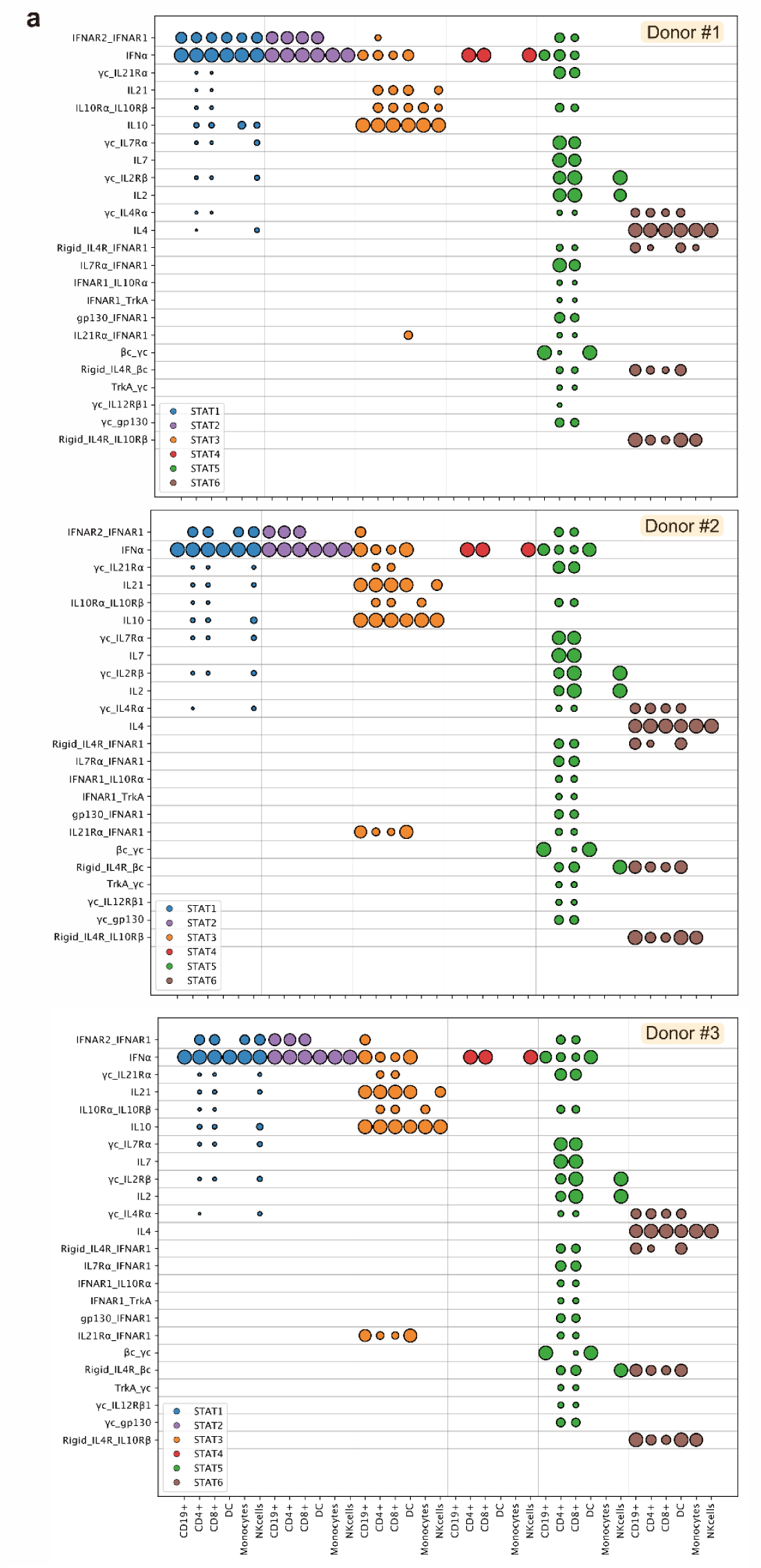
Agonists activity across PBMC cell populations. **(a)** cell-specific activity of selected novel agonists in the pSTAT1, pSTAT2, pSTAT3, pSTAT4, pSTAT5 and pSTAT6 pathways across PBMC cell subtypes from the three donors tested.

**Supplementary Figure 7:**
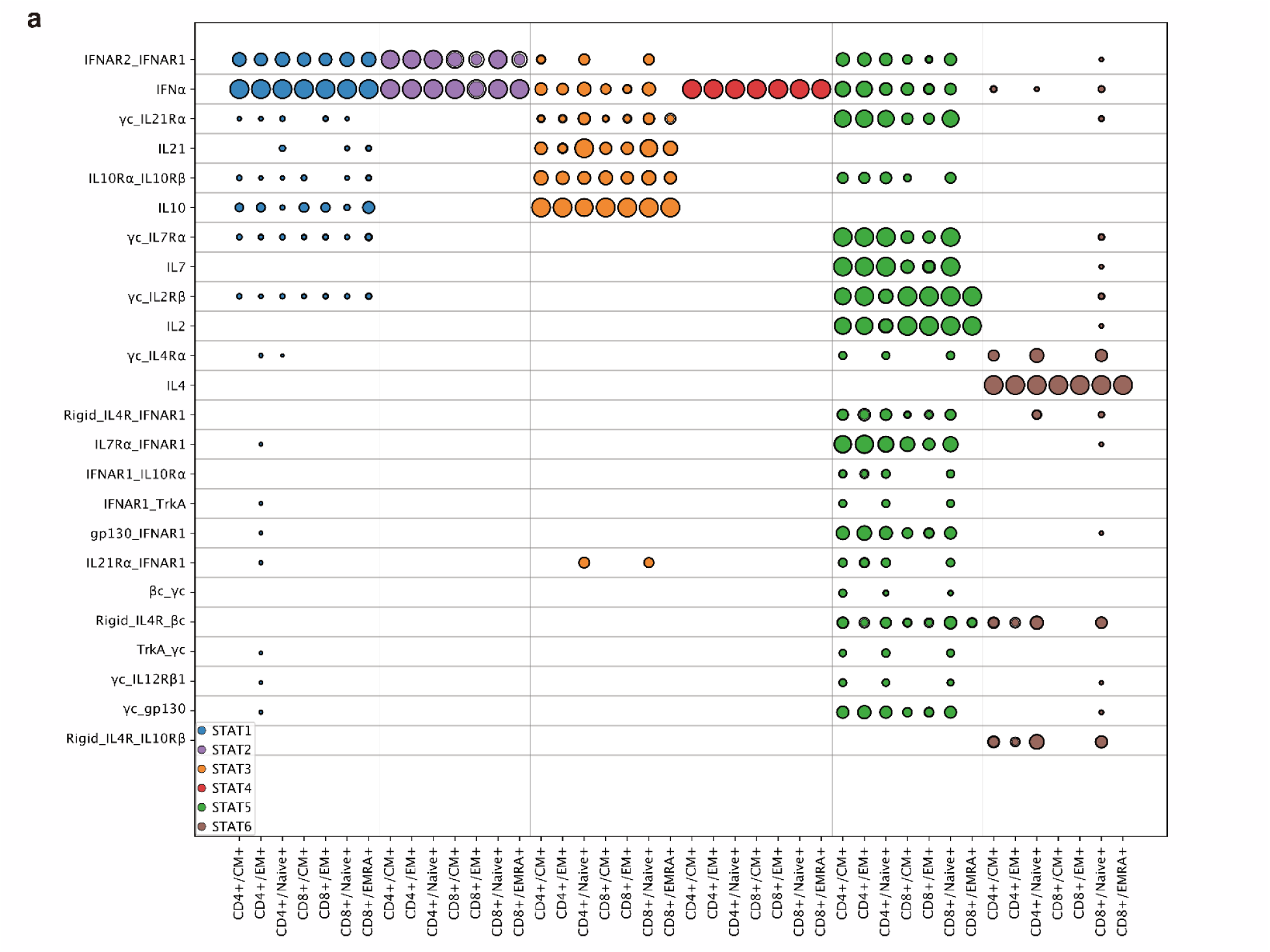
Agonists activity across T-cell subsets. **(a)** cell-specific activity of selected novel agonists in the pSTAT1, pSTAT2, pSTAT3, pSTAT4, pSTAT5 and pSTAT6 pathways across T-cell subtypes. Results displayed are averages from the three donors tested.

**Supplementary Table 1:**
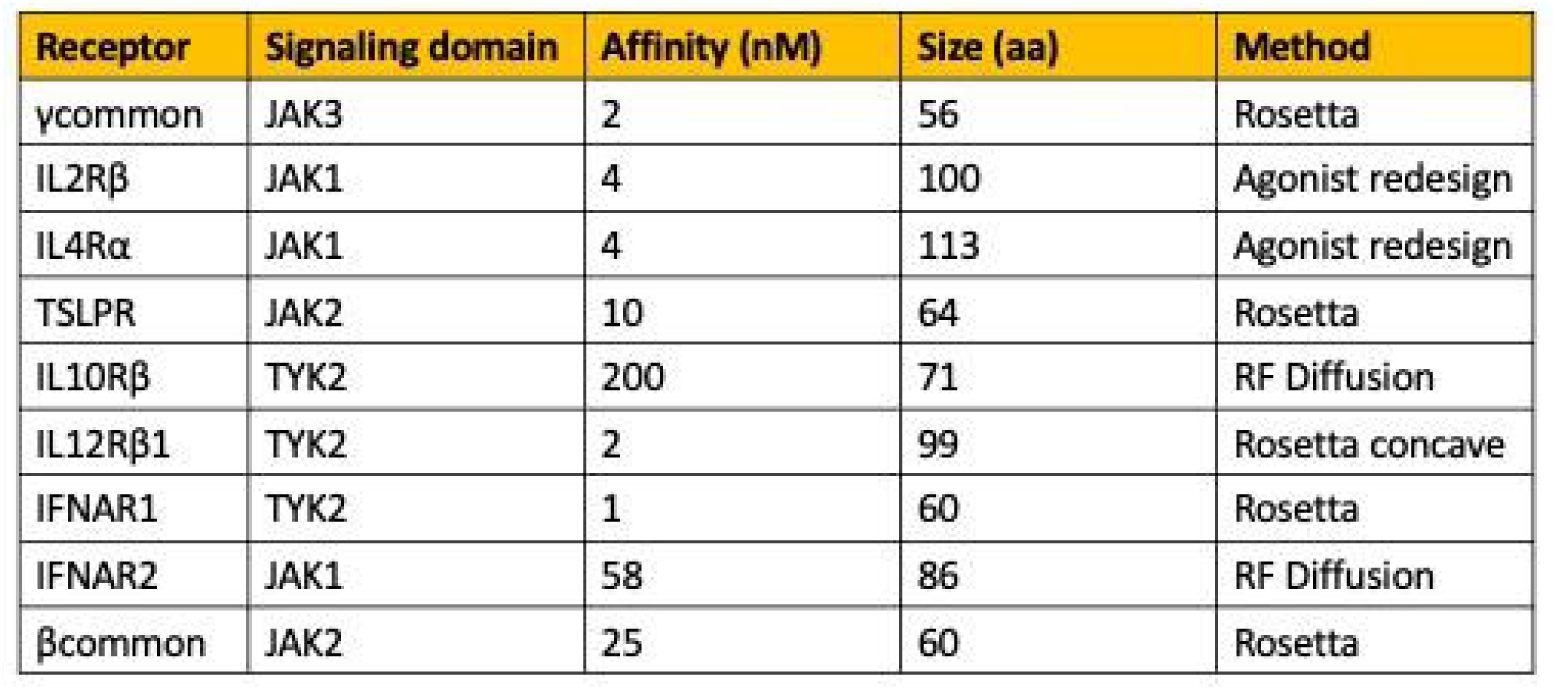
List of receptor binding proteins generated in this study. (a) summary table detailing the receptor binding proteins generated in this study. The table includes information on the target receptor, signaling domain, affinity, size, and design method for each binder.

**Supplementary Table 2:**
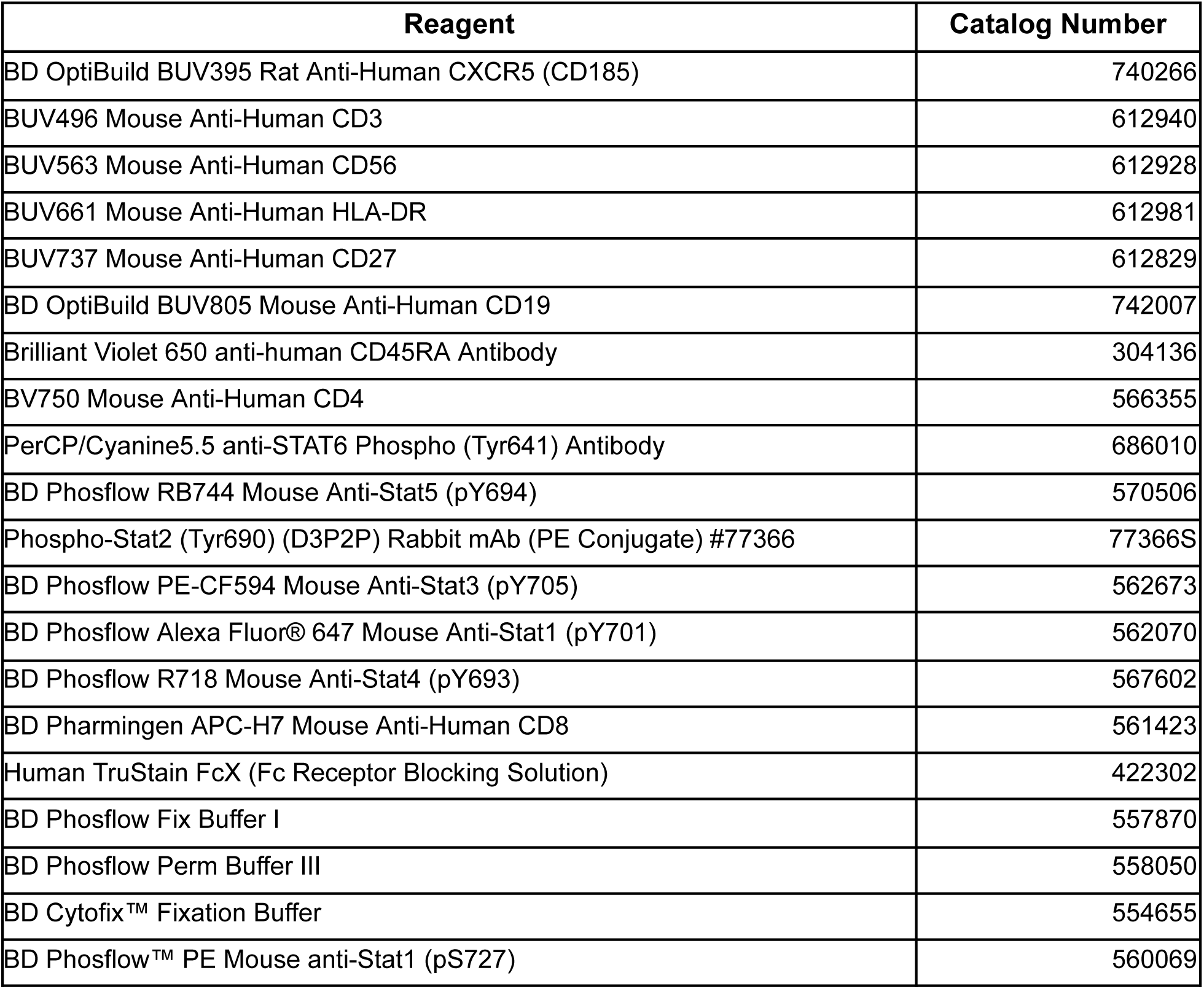
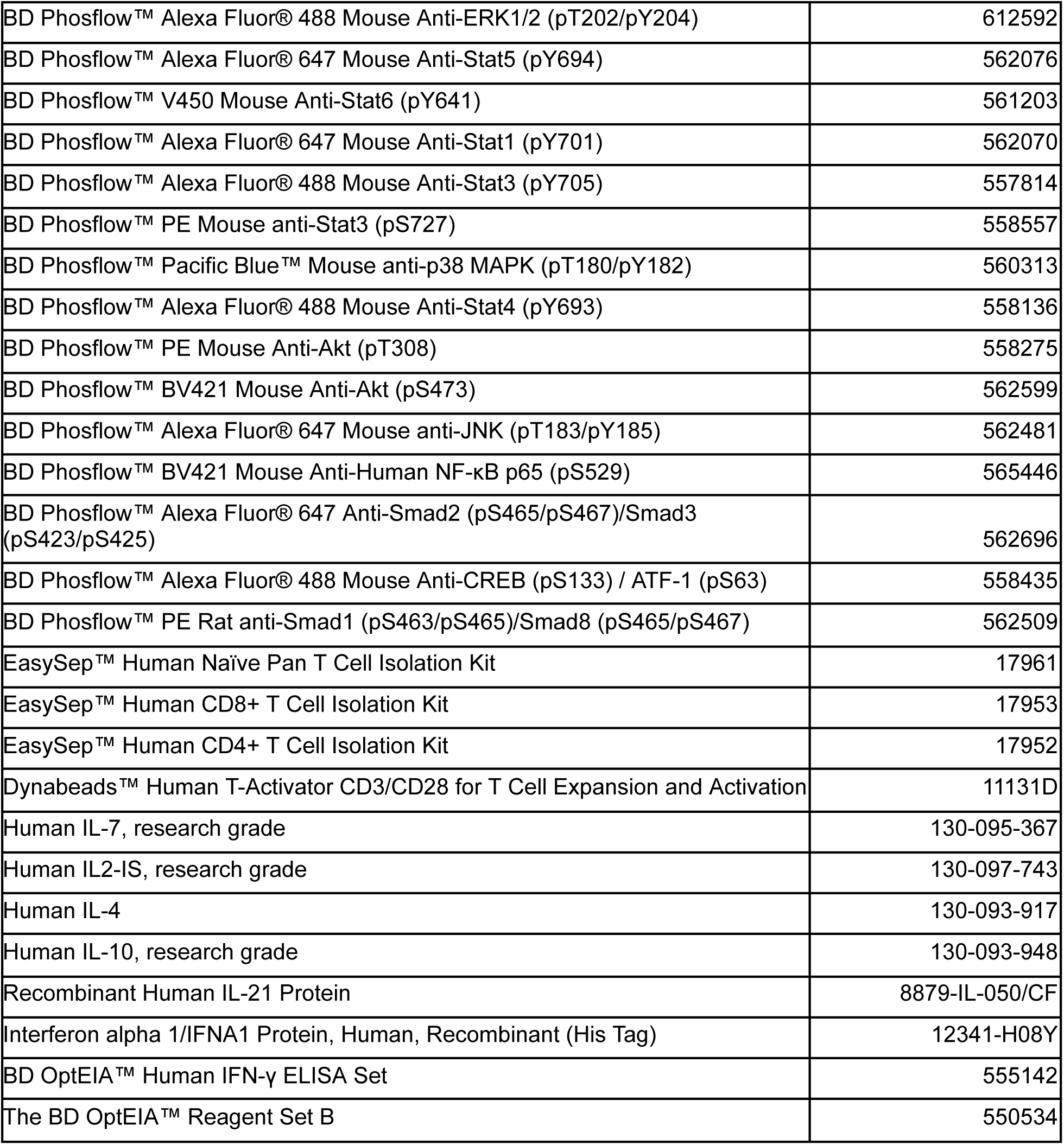
List of reagents used in this study.

## Methods

### Binder Design

Binders against human receptors were developed using three methodologies: Rosetta-based methods, RFdiffusion, and redesign of agonists. For the Rosetta-based approach, we followed the framework described in Cao et al^13^, utilizing Rosetta’s physical principles to select residues that target the desired interface and identifying suitable scaffolds from our libraries. The RFdiffusion method employs a deep learning model to generate protein backbones through denoising with an optimized loss function that targets specific "hot spots" for binding. We then applied a Message Passing Neural Network (MPNN)^42^ to optimize the sequence corresponding to the generated backbone, as RFdiffusion produces backbones without sequences. The final sequences were predicted using AlphaFold2, filtered by the pLDDT (a metric that predicts confidence in the sequence forming the desired structure) and PAE metrics for binding likelihood. Lastly, in the redesign of cytokines, we fixed the binding interface residues on one side while using MPNN to modify the opposing side’s binding interface, adapting it to a new context where it is exposed to water without a receptor. In this way we maintained one binding interface while mitigating the other.

### Experimental Screening of Binders

Protein binder designs were synthesized as synthetic genes (eBlocks, Integrated DNA Technologies) with compatible BsaI overhangs for cloning into the modified expression vector LM0627, which contains a Kanamycin resistance gene and a ccdb lethal gene. Subcloning produced the following product: MSG-[protein]GSGSHHWGSTHHHHHH, featuring a C-terminal SNAC cleavage tag and a 6XHis affinity tag. Helical peptide binders were ordered in a similar format, with added adaptors (GGGSGGGGSASHMRS, SSEISFCSEPPPSRRS) that allowed cloning into both the pETcon3 vector and LM0627, facilitating purification in E. coli and yeast surface display.

For protein binder expression screens, we followed a previously reported protocol^20^, with some modifications. Golden Gate subcloning reactions were conducted in 96-well PCR plates at a volume of 1 µL. The reaction mixtures were transformed into a chemically competent expression strain (BL21(DE3)), followed by 1-hour outgrowths, which were split into four 96-deep well plates containing 0.9-1.0 mL of auto-induction media (autoclaved TBII media supplemented with Kanamycin, 2 mM MgSO4, 1X 5052) for a final volume of approximately 4 mL. After 20-24 hours, cells were harvested and lysed, and clarified lysates were applied to a 50 µL bed of Ni-NTA agarose resin in a 96-well fritted plate equilibrated with a Tris wash buffer. Following sample application and flow-through, the resin was thoroughly washed, and samples were eluted in 200 µL of a Tris elution buffer containing 300 mM imidazole. All eluates were sterile filtered with a 96-well 0.22 µm filter plate (Agilent 203940-100) prior to size exclusion chromatography.

Protein designs were screened via SEC using an AKTA FPLC outfitted with an autosampler capable of running samples from a 96-well source plate. The protein binders were run on a Superdex 75 Increase 5/150 GL column (Cytiva 29148722). To improve peak resolution, the SEC column was connected directly in line from the autosampler to the UV detector. 0.25 mL fractions were collected from each run, and selected fractions were pooled for further analysis (LC-MS, native mass spectrometry, negative stain EM, SDS-PAGE).

Binding affinity to target receptors was measured using a Biacore 8K with a Protein A chip (Cytiva #29127556) at a target concentration of 5 µg/mL. Binding traces were collected by running single-cycle kinetics at increased binder concentrations (four series with 8-fold titration starting at 1000 nM), with binders and targets diluted in HEPES buffer (Cytiva #BR100669).

### Protein Expression of agonists

In addition to the standard purification of protein binders described above, two additional steps were performed for agonist protein expression. The insoluble fraction was resuspended in 6 M guanidine hydrochloride and applied to the Ni-NTA resin. The concentration of guanidine was gradually reduced to refold the protein, with the soluble fraction added on top. Additionally cells underwent low endotoxin steps where proteins were washed with 0.7% CHAPS buffer four times after incubating for 10 minutes at each step. All eluates were sterile filtered with a 96-well 0.22 µm filter plate (Agilent 203940-100) prior to size exclusion chromatography.

### Cell Lines and PBMCs

For this study, the following cell lines were used: wild-type Nalm6, TF-1, and U937 (ATCC). All cell lines were cultured in RPMI-1640 Medium with 1% GlutaMAX (Gibco Cat# 61870036), supplemented with 10% fetal bovine serum (Life Technologies Cat# A4766801) and 1% penicillin-streptomycin (Gibco Cat# 15140122). TF-1 cells were additionally supplemented with 2 ng/mL of recombinant human GM-CSF (Biotechne Cat# 7954-Gm) to support their growth. The cells were passed every 3 days to avoid over-confluency and maintained at 37°C in a humidified incubator with 5% CO2. Peripheral Blood Mononuclear Cells (PBMCs) were obtained from Bloodworks Seattle, thawed, washed with DNase, and either rested for one hour before experimentation or subjected to a 3-hour starvation period prior to assays if required.

### Cell Signaling Assays

Cell signaling assays were conducted according to a unified protocol. Cells were initially stimulated with the relevant ligand for 15 minutes. Following stimulation, cells were fixed using BD Cytofix Fixative (formaldehyde), permeabilized, and then stained with antibodies detailed in Supplementary Table 2. Signal quantification was performed via flow cytometry. For inhibition assays, cells were pre-incubated with the binders for 30 minutes before the introduction of the cytokine intended for blockade. After cytokine addition, cells were further stimulated for 15 minutes.

### Receptor Knockout

Cell lines for knockout tests were first selected by confirming that the designed agonists and the appropriate natural cytokine controls signaled as expected in the wild-type background using phospho-flow cytometry, as described above. Single receptor knockouts were generated by nucleofection with multi-guide sgRNAs (Synthego Gene Knockout Kit v2) according to the manufacturer’s instructions. Briefly, multi-guide sgRNAs targeting the genes of interest were purchased (Synthego), diluted to 30 µM in the appropriate nucleofection buffer (SE buffer, Lonza Cat# V4XC-1032; or SF buffer, Lonza Cat# V4XC-2032), and combined with Cas9 to form ribonucleoprotein (RNP) complexes. After a 10-minute incubation, 2.0 × 10^5 cells were added to the pre-complexed RNPs and electroporated using a 4D-X Core unit (Lonza Cat# AAF-1003B). Cells were recovered in 2 mL pre-warmed RPMI media and expanded over one week for downstream assays. Knockouts were confirmed by antibody staining against the receptors of interest and compared to isotype controls. If >10% of cells remained positively stained, the receptor-negative population was enriched by cell sorting (BD FACSAria II) and scaled up. In some cases where electroporation did not yield complete knockout, single-cell clones were isolated in 384-well plates and individually screened for loss of signaling. Receptor validation was performed using natural cytokine responsiveness or novokine signaling, depending on the receptor. IFNAR1 knockouts were confirmed by loss of response to IFNα. Because U937 cells do not express IL7Rα, IL7Rα was stably introduced by lentiviral transduction, and successful overexpression was verified by restored responsiveness to IL7. IL21Rα knockouts were validated by loss of response to IL21 cytokine, while γcommon knockouts were confirmed by loss of responsiveness to IL21. For βcommon and gp130, where parental U937 cells do not respond to their natural cytokines, knockout validation was assessed by loss of signaling in response to the designed novokines.

### Design of rigid agonists

Rigid versions of the agonists were designed following the protocol previously descrived^40^. Briefly, the binder-receptor structures were positioned next to each other for each of the novokines to design. Then, thousands of geometries for each pair were sampled and filtered for clashes using PyMOL, and the two binders were fused into a rigid, single-chain protein using RFdiffusion setting the binders as contigs and giving a variable range of length for the scaffold. The sequence of these scaffolds was then designed using ProteinMPNN excluding the binder interfaces to preserve binding, and filtered using Alphafold2 to check for agreement between the predicted and designed models. Designs were selected for experimental testing based on low RMSD, high pLDDT (Alphafold2 confidence in the prediction) and diversity of induced receptor geometries.

### Profiling signaling signature and cell specificity in PBMCs

PBMCs were thawed and rested for 1 hour at 37 °C in complete RPMI supplemented with 5% FBS, 1× monensin, and 1× brefeldin A (BFA). Following the rest period, PBMCs were stimulated with a final concentration of 250 nM of either natural or synthetic cytokines for 15 minutes at 37°C. Reactions were stopped by fixation with BD Phosflow Fix Buffer I for 15 minutes at 37 °C. Cells were then pelleted and resuspended in 50 µL of staining buffer containing surface lineage markers. After 30 minutes, cells were washed, resuspended in 100 µL of BD Phosflow Perm Buffer III, and incubated on ice for 1 hour. Following permeabilization, cells were washed and stained overnight at 4C with a master mix containing additional lineage markers and antibodies specific for phosphorylated STAT1-6 (pSTAT1-6). The following day, cells were resuspended in the staining buffer, and data were acquired on a Cytek Aurora flow cytometer. For analysis, signaling was normalized per patient, cell type, and pSTAT on a 0-1 scale. The mean and standard error of the mean (SEM) were calculated from three unstimulated controls. A condition was considered signaling if it exceeded the mean by at least 6xSEM. For samples with weak positive cytokine responses, the SEM cutoff was increased to prevent noise from being misclassified as signal.

### Monocyte proliferation assay

TF-1 cells were starved overnight in RPMI medium with 10% FBS but without GM-CSF supplementation. For the proliferation assay, 15k cells/well were stimulated with 250nM of the agonists in a 384 well plate, and proliferation was measured after 72h by flow cytometry (Attune NxT Flow Cytometer).

### T cell survival assay

Naive T cells were isolated using a StemCell Technologies kit and subsequently distributed into a 384-well plate. Cells were incubated with the designed ligands, which were refreshed every 3 days. Cell counts were measured after 8 days to assess T cell survival.

### Proliferation of T Cells Assay

CD4 and CD8 T cells were isolated from PBMCs using a StemCell Technologies kit. Purified pan T cells were then incubated with CD3/CD28 beads for 2 days and then rested for no beads for one day. On the fourth day, the beads were removed, and the cells were distributed into a 384-well plate. Designed ligands were added to each well in triplicate and replenished every 4 days. Cells were diluted based on their confluence state. Cells were counted using flow cytometry to quantify their proliferation.

### IFNγ Secretion Assay

CD8 T cells were isolated from PBMCs using a StemCell Technologies isolation kit. Purified CD8 T cells were expanded with CD3/CD28 Dynabeads and IL-2 for 3 days, after which the beads were removed. Cells were further expanded in the presence of IL-2 with fresh cytokine replenishment. Once expansion slowed, cells were deprived of IL-2 for 2 days before being plated into 384-well plates in triplicate. Cells were cultured with novokines or control cytokines for 3 days to prime them. On day 4, cells were stimulated with anti-CD3 antibodies for 4 hours to trigger IFNγ secretion. Following stimulation, culture media was collected, and secreted IFNγ levels were quantified using ELISA.

## References

1. Akdis, M., Burgler, S., Crameri, R., Eiwegger, T., Fujita, H., Gomez, E., Klunker, S., Meyer, N., O’Mahony, L., Palomares, O., et al. (2011). Interleukins, from 1 to 37, and interferon-γ: Receptors, functions, and roles in diseases. J. Allergy Clin. Immunol. 127, 701–721.e70. 10.1016/j.jaci.2010.11.050.

2. Cui, A., Huang, T., Li, S., Ma, A., Pérez, J.L., Sander, C., Keskin, D.B., Wu, C.J., Fraenkel, E., and Hacohen, N. (2024). Dictionary of immune responses to cytokines at single-cell resolution. Nature 625, 377–384. 10.1038/s41586-023-06816-9.

3. Pires, I.S., Hammond, P.T., and Irvine, D.J. (2021). Engineering Strategies for Immunomodulatory Cytokine Therapies: Challenges and Clinical Progress. Adv. Ther. 4, 2100035. 10.1002/adtp.202100035.

4. Moraga, I., Spangler, J.B., Mendoza, J.L., Gakovic, M., Wehrman, T.S., Krutzik, P., and Garcia, K.C. Synthekines are surrogate cytokine and growth factor agonists that compel signaling through non-natural receptor dimers. eLife 6, e22882. 10.7554/eLife.22882.

5. Yen, M., Ren, J., Liu, Q., Glassman, C.R., Sheahan, T.P., Picton, L.K., Moreira, F.R., Rustagi, A., Jude, K.M., Zhao, X., et al. (2022). Facile discovery of surrogate cytokine agonists. Cell 185, 1414–1430.e19. 10.1016/j.cell.2022.02.025.

6. Doiron, K.E., Way, J.C., and Silver, P.A. (2025). Rational design of selective bispecific EPO-R/CD131 agonists. Preprint at bioRxiv, 10.1101/2025.04.10.648227 https://doi.org/10.1101/2025.04.10.648227.

7. Doiron, K.E., Way, J.C., and Silver, P.A. (2025). Rational design of selective bispecific EPO-R/CD131 agonists. Preprint at bioRxiv, 10.1101/2025.04.10.648227 https://doi.org/10.1101/2025.04.10.648227.

8. Zhao, Y., Ogishi, M., Pal, A., Su, L.L., Tao, P., Jiang, H., Rodriguez, G.E., Chen, X., Sun, Q., Rysavy, L.W., et al. (2025). Expanding the cytokine receptor alphabet reprograms T cells into diverse states. Nature, 1–12. 10.1038/s41586-025-09393-1.

9. Mohan, K., Ueda, G., Kim, A.R., Jude, K.M., Fallas, J.A., Guo, Y., Hafer, M., Miao, Y., Saxton, R.A., Piehler, J., et al. (2019). Topological control of cytokine receptor signaling induces differential effects in hematopoiesis. Science 364, eaav7532. 10.1126/science.aav7532.

10. Chun, J.-H., Lim, B.S., Roy, S., Walsh, M.J., Abhiraman, G.C., Zhangxu, K., Atajanova, T., Revach, O.-Y., Clark, E.C., Li, P., et al. (2024). Potent antitumor activity of a designed interleukin-21 mimic. Preprint at bioRxiv, 10.1101/2024.12.06.626481 https://doi.org/10.1101/2024.12.06.626481.

11. Yang, H., Ulge, U.Y., Quijano-Rubio, A., Bernstein, Z.J., Maestas, D.R., Chun, J.-H., Wang, W., Lin, J.-X., Jude, K.M., Singh, S., et al. (2023). Design of cell-type-specific hyperstable IL-4 mimetics via modular de novo scaffolds. Nat. Chem. Biol. 19, 1127–1137. 10.1038/s41589-023-01313-6.

12. Silva, D.-A., Yu, S., Ulge, U.Y., Spangler, J.B., Jude, K.M., Labão-Almeida, C., Ali, L.R., Quijano-Rubio, A., Ruterbusch, M., Leung, I., et al. (2019). De novo design of potent and selective mimics of IL-2 and IL-15. Nature 565, 186–191. 10.1038/s41586-018-0830-7.

13. Cao, L., Coventry, B., Goreshnik, I., Huang, B., Sheffler, W., Park, J.S., Jude, K.M., Marković, I., Kadam, R.U., Verschueren, K.H.G., et al. (2022). Design of protein-binding proteins from the target structure alone. Nature 605, 551–560. 10.1038/s41586-022-04654-9.

14. Yang, W., Hicks, D.R., Ghosh, A., Schwartze, T.A., Conventry, B., Goreshnik, I., Allen, A., Halabiya, S.F., Kim, C.J., Hinck, C.S., et al. (2024). Design of High Affinity Binders to Convex Protein Target Sites. Preprint at bioRxiv, 10.1101/2024.05.01.592114 https://doi.org/10.1101/2024.05.01.592114.

15. Watson, J.L., Juergens, D., Bennett, N.R., Trippe, B.L., Yim, J., Eisenach, H.E., Ahern, W., Borst, A.J., Ragotte, R.J., Milles, L.F., et al. (2023). De novo design of protein structure and function with RFdiffusion. Nature 620, 1089–1100. 10.1038/s41586-023-06415-8.

16. Hunt, A.C., Case, J.B., Park, Y.-J., Cao, L., Wu, K., Walls, A.C., Liu, Z., Bowen, J.E., Yeh, H.-W., Saini, S., et al. (2022). Multivalent designed proteins neutralize SARS-CoV-2 variants of concern and confer protection against infection in mice. Sci. Transl. Med. 14, eabn1252. 10.1126/scitranslmed.abn1252.

17. Ebina-Shibuya, R., and Leonard, W.J. (2023). Role of thymic stromal lymphopoietin in allergy and beyond. Nat. Rev. Immunol. 23, 24–37. 10.1038/s41577-022-00735-y.

18. Berger, S., Seeger, F., Yu, T.-Y., Aydin, M., Yang, H., Rosenblum, D., Guenin-Macé, L., Glassman, C., Arguinchona, L., Sniezek, C., et al. (2024). Preclinical proof of principle for orally delivered Th17 antagonist miniproteins. Cell 187, 4305–4317.e18. 10.1016/j.cell.2024.05.052.

19. Morris, R.M., Mortimer, T.O., and O’Neill, K.L. (2022). Cytokines: Can Cancer Get the Message? Cancers 14, 2178. 10.3390/cancers14092178.

20. Sappington, I., Toul, M., Lee, D.S., Robinson, S.A., Goreshnik, I., McCurdy, C., Chan, T.C., Buchholz, N., Huang, B., Vafeados, D., et al. (2024). Improved protein binder design using beta-pairing targeted RFdiffusion. Preprint at bioRxiv, 10.1101/2024.10.11.617496 https://doi.org/10.1101/2024.10.11.617496.

21. Glögl, M., Krishnakumar, A., Ragotte, R.J., Goreshnik, I., Coventry, B., Bera, A.K., Kang, A., Joyce, E., Ahn, G., Huang, B., et al. (2024). Target-conditioned diffusion generates potent TNFR superfamily antagonists and agonists. Science 386, 1154–1161. 10.1126/science.adp1779.

22. Wang, X., Cardoso, S., Cai, K., Venkatesh, P., Hung, A., Ng, M., Hall, C., Coventry, B., Lee, D., Chowhan, R., et al. (2024). Tuning Insulin Receptor Signaling Using De Novo Designed Agonists. Preprint at bioRxiv, 10.1101/2024.10.07.617068 https://doi.org/10.1101/2024.10.07.617068.

23. Huang, B., Abedi, M., Ahn, G., Coventry, B., Sappington, I., Tang, C., Wang, R., Schlichthaerle, T., Zhang, J.Z., Wang, Y., et al. (2025). Designed endocytosis-inducing proteins degrade targets and amplify signals. Nature 638, 796–804. 10.1038/s41586-024-07948-2.

24. Huang, B., Coventry, B., Borowska, M.T., Arhontoulis, D.C., Exposit, M., Abedi, M., Jude, K.M., Halabiya, S.F., Allen, A., Cordray, C., et al. (2024). De novo design of miniprotein antagonists of cytokine storm inducers. Nat. Commun. 15, 7064. 10.1038/s41467-024-50919-4.

25. Woldman, I., Varinou, L., Ramsauer, K., Rapp, B., and Decker, T. (2001). The Stat1 Binding Motif of the Interferon-γ Receptor Is Sufficient to Mediate Stat5 Activation and Its Repression by SOCS3 *. J. Biol. Chem. 276, 45722–45728. 10.1074/jbc.M105320200.

26. May, P., Gerhartz, C., Heesel, B., Welte, T., Doppler, W., Graeve, L., Horn, F., and Heinrich, P.C. (1996). Comparative study on the phosphotyrosine motifs of different cytokine receptors involved in STAT5 activation. FEBS Lett. 394, 221–226. 10.1016/0014-5793(96)00955-6.

27. Hanna, B.S., Llaó-Cid, L., Iskar, M., Roessner, P.M., Klett, L.C., Wong, J.K.L., Paul, Y., Ioannou, N., Öztürk, S., Mack, N., et al. (2021). Interleukin-10 receptor signaling promotes the maintenance of a PD-1int TCF-1+ CD8+ T cell population that sustains anti-tumor immunity. Immunity 54, 2825–2841.e10. 10.1016/j.immuni.2021.11.004.

28. Cui, W., Liu, Y., Weinstein, J.S., Craft, J., and Kaech, S.M. (2011). An Interleukin-21- Interleukin-10-STAT3 Pathway Is Critical for Functional Maturation of Memory CD8+ T Cells. Immunity 35, 792–805. 10.1016/j.immuni.2011.09.017.

29. Jahn, T., Sindhu, S., Gooch, S., Seipel, P., Lavori, P., Leifheit, E., and Weinberg, K. (2007). Direct interaction between Kit and the interleukin-7 receptor. Blood 110, 1840–1847. 10.1182/blood-2005-12-028019.

30. Wang, Y., van Boxel-Dezaire, A.H.H., Cheon, H., Yang, J., and Stark, G.R. (2013). STAT3 activation in response to IL-6 is prolonged by the binding of IL-6 receptor to EGF receptor. Proc. Natl. Acad. Sci. 110, 16975–16980. 10.1073/pnas.1315862110.

31. Hardbower, D.M., Singh, K., Asim, M., Verriere, T.G., Olivares-Villagómez, D., Barry, D.P., Allaman, M.M., Washington, M.K., Peek, R.M., Piazuelo, M.B., et al. EGFR regulates macrophage activation and function in bacterial infection. J. Clin. Invest. 126, 3296–3312. 10.1172/JCI83585.

32. MacDonald, F., and Zaiss, D.M.W. (2017). The Immune System’s Contribution to the Clinical Efficacy of EGFR Antagonist Treatment. Front. Pharmacol. 8. 10.3389/fphar.2017.00575.

33. Zeboudj, L., Maître, M., Guyonnet, L., Laurans, L., Joffre, J., Lemarie, J., Bourcier, S., Nour-Eldine, W., Guérin, C., Friard, J., et al. (2018). Selective EGF-Receptor Inhibition in CD4+ T Cells Induces Anergy and Limits Atherosclerosis. J. Am. Coll. Cardiol. 71, 160–172. 10.1016/j.jacc.2017.10.084.

34. Huang, L., Zhang, X., Fan, J., Liu, X., Luo, S., Cao, D., Liu, Y., Xia, Z., Zhong, H., Chen, C., et al. (2023). EGFR promotes the apoptosis of CD4+ T lymphocytes through TBK1/Glut1 induced Warburg effect in sepsis. J. Adv. Res. 44, 39–51. 10.1016/j.jare.2022.04.010.

35. Minutti, C.M., Drube, S., Blair, N., Schwartz, C., McCrae, J.C., McKenzie, A.N., Kamradt, T., Mokry, M., Coffer, P.J., Sibilia, M., et al. (2017). Epidermal Growth Factor Receptor Expression Licenses Type-2 Helper T Cells to Function in a T Cell Receptor-Independent Fashion. Immunity 47, 710–722.e6. 10.1016/j.immuni.2017.09.013.

36. Ley, S., Weigert, A., Weichand, B., Henke, N., Mille-Baker, B., Janssen, R. a. J., and Brüne, B. (2013). The role of TRKA signaling in IL-10 production by apoptotic tumor cell-activated macrophages. Oncogene 32, 631–640. 10.1038/onc.2012.77.

37. Lambiase, A., Bracci-Laudiero, L., Bonini, S., Bonini, S., Starace, G., D’Elios, M.M., Carli, M.D., and Aloe, L. (1997). Human CD4+ T cell clones produce and release nerve growth factor and express high-affinity nerve growth factor receptors. J. Allergy Clin. Immunol. 100, 408–414. 10.1016/S0091-6749(97)70256-2.

38. Yin, T., Wang, G., Wang, L., Mudgal, P., Wang, E., Pan, C.C., Alexander, P.B., Wu, H., Cao, C., Liang, Y., et al. (2024). Breaking NGF–TrkA immunosuppression in melanoma sensitizes immunotherapy for durable memory T cell protection. Nat. Immunol. 25, 268–281. 10.1038/s41590-023-01723-7.

39. Kalie, E., Jaitin, D.A., Podoplelova, Y., Piehler, J., and Schreiber, G. (2008). The Stability of the Ternary Interferon-Receptor Complex Rather than the Affinity to the Individual Subunits Dictates Differential Biological Activities*. J. Biol. Chem. 283, 32925–32936. 10.1074/jbc.M806019200.

40. Marc Expòsit, Mohamad Abedi, Aditya Krishnakumar, and David Baker (09252025). Geometric Tuning of Cytokine Receptor Association Modulates Synthetic Agonist Signaling. Preprint at bioRxiv.

41. Engelowski, E., Schneider, A., Franke, M., Xu, H., Clemen, R., Lang, A., Baran, P., Binsch, C., Knebel, B., Al-Hasani, H., et al. (2018). Synthetic cytokine receptors transmit biological signals using artificial ligands. Nat. Commun. 9, 2034. 10.1038/s41467-018-04454-8.

42. Dauparas, J., Anishchenko, I., Bennett, N., Bai, H., Ragotte, R.J., Milles, L.F., Wicky, B.I.M., Courbet, A., de Haas, R.J., Bethel, N., et al. (2022). Robust deep learning–based protein sequence design using ProteinMPNN. Science 378, 49–56. 10.1126/science.add2187.

